# The RIDD activity of *C. elegans* IRE1 modifies neuroendocrine signaling in anticipation of environment stress to ensure survival

**DOI:** 10.1101/2023.08.10.552841

**Authors:** Mingjie Ying, Yair Argon, Tali Gidalevitz

## Abstract

*Xbp1* splicing and regulated IRE1-dependent RNA decay (RIDD) are two RNase activities of the ER stress sensor IRE1. While *Xbp1* splicing has important roles in stress responses and animal physiology, the physiological role(s) of RIDD remain enigmatic. Genetic evidence in *C. elegans* connects XBP1-independent IRE1 activity to organismal stress adaptation, but whether this is via RIDD, and what are the targets is yet unknown. We show that cytosolic kinase/RNase domain of *C. elegans* IRE1 is indeed capable of RIDD in human cells, and that sensory neurons use RIDD to signal environmental stress, by degrading mRNA of TGFβ-like growth factor DAF-7. *daf-7* was degraded in human cells by both human and worm IRE1 RNAse activity with same efficiency and specificity as *Blos1,* confirming *daf-7* as RIDD substrate. Surprisingly, *daf-7* degradation *in vivo* was triggered by concentrations of ER stressor tunicamycin too low for *xbp-1* splicing. Decrease in DAF-7 normally signals food limitation and harsh environment, triggering adaptive changes to promote population survival. Because *C. elegans* is a bacteriovore, and tunicamycin, like other common ER stressors, is an antibiotic secreted by *Streptomyces spp.*, we asked whether *daf-7* degradation by RIDD could signal pending food deprivation. Indeed, pre-emptive tunicamycin exposure increased survival of *C. elegans* populations under food limiting/high temperature stress, and this protection was abrogated by overexpression of DAF-7. Thus, *C. elegans* uses stress-inducing metabolites in its environment as danger signals, and employs IRE1’s RIDD activity to modulate the neuroendocrine signaling for survival of upcoming environmental challenge.

## Introduction

Serine/Threonine-Protein Kinase/Endoribonuclease IRE1 (inositol-requiring enzyme 1) is the most ancient and conserved sensor in the unfolded protein response (UPR) pathway. Its main function is to resolve proteostatic stress in the endoplasmic reticulum (ER), or, alternatively, trigger apoptosis when encountering prolonged and severe stress^1^. IRE1 is a single spanning type I trans-membrane protein^2^ with a stress-sensing lumenal domain and an unusual combined kinase and RNase domain located in cytosol^3,4^. Upon ER stress, IRE1 undergoes dimerization and oligomerization^5,6^, accompanied by a series of conformational changes that initiate from the lumenal domain and proceed to the active conformation of the RNase domain^3,7–9^. Activated IRE1 has two distinct RNase activities: 1) cleaving of mRNA of the X-box binding protein (XBP1) transcription factor^10–12^, resulting in removal of a small non-canonical intron and, upon ligation by the tRNA ligase RTCB^13,14^, production of an active transcription factor *Xbp1*s (known as ‘splicing activity’), and 2) cleaving a set of transcripts and microRNAs, resulting in their degradation by exonucleases (known as the regulated IRE1 dependent decay of RNA, or RIDD)^15^. The *Xbp1* splicing activity of IRE1 is strongly conserved across evolution, with the fission yeast *Schizosaccharomyces pombe* being the only identified exception so far^16^. Importantly, in addition to orchestrating the transcriptional response to acute stress in the ER, the *Xbp1* splicing activity functions in multiple aspects of metazoan physiology, such as differentiation of secretory cell types, or lipid metabolism^17–21^.

In contrast to the splicing activity, the functional importance of IRE1’s RIDD activity remains poorly understood. In cells, activation of RIDD by severe stress has been associated with either pro- or anti-apoptotic outcomes via degradation of specific RIDD targets. For example, RIDD activity increases caspase-2 synthesis by degrading micro RNAs^22^, but inhibits caspase-8 activation by promoting decay of DR-5^23^. Many findings also indicate that RIDD helps maintain protein homeostasis in the ER under acute stress, but it does so by different means in different cell types and organisms. For example, in fission yeast *S. pombe*^16^ and in *Drosophila* S2 cells^24^, IRE1 RIDD activity degrades a broad range of mRNAs, decreasing the biosynthetic load in the ER. In contrast, in mammalian cells, this protection mechanism seems more restricted, since both the number of targets and the magnitude of downregulation are lower than in yeast and *Drosophila* cells^24–26^. Instead, RIDD protects these cells from ER stress by degrading individual targets. For example, degradation of the 28S rRNA by the human IRE1b decreases protein synthesis under ER stress in intestinal cells^27^; similarly, translational attenuation may result from RIDD of mRNA for a regulatory subunit of eukaryotic translation initiation factor 2α (eIF2α) phosphatase, by IRE1α^28^; third, degradation of the *Blos1* transcript may assist the clearance of misfolded species and aggregates by reorganizing early endosomes and lysosomes^29,30^.

At organismal level, ectopic activation of RIDD, either by acute ER stress or by deleting XBP1 (which results in compensatory over-activation of IRE1^31^), can also lead to both protective or detrimental outcomes^32^. In *C. elegans*, IRE-1-dependent degradation of *flp-6* mRNA, coding for a neuropeptide secreted by the ASE sensory neuron, resulted in abnormal somatic differentiation of germ cells in animals genetically prone to forming germline tumors^33^. In mice, RIDD may contribute to neuroinflammation under hypoxia–ischemia stress by targeting miR-17^34^, which in turn activates NLRP3 inflammasome^35^. On the other hand, activation of RIDD in the liver protected mice from hepatotoxicity^36,37^.

The physiological *in vivo* functions of RIDD are even less understood, for three reasons. First, it has been challenging to identify *in vivo* RIDD targets. Unlike for *Xbp1*, there is no definitive mRNA structure of a RIDD target that specifies IRE1 recognition and cleavage. The *Xbp1*-like stem-loops have only been found in some RIDD substrates, including the most canonical substrate *Blos1*^25,38^. The other RIDD-targeted transcripts have no *Xbp1* like stem-loops or cleavage sites, yet are downregulated in stress-dependent manner by IRE1^22,39,40^. Moreover, none of the 13 *in vitro* - identified RIDD targets that were used to define the consensus recognition/cleavage sequence showed ER stress-dependent decay in the same study^41^. Thus, identification of *in vivo* RIDD targets mainly relies on analyzing the transcriptomic changes in IRE1- and ER stress-dependent manner.

Second, it is difficult to uncouple RIDD from *Xbp1* splicing mechanistically, or to experimentally activate RIDD without affecting *Xbp1* splicing. Yet, RIDD is selectively and physiologically activated in specialized secretory cells that experience high biosynthetic flux, such as insulin-producing pancreatic beta cells, mucin-producing goblet cells, and chylomicrons-producing enterocytes, where it limits the ER overload by targeting the messages of high-flux proteins^42–45^. That RIDD is protective in these cells indicates that its selective physiological activation may serve as an adaptive strategy.

Finally, there are few identified examples to date where a physiological *in vivo* function of RIDD is not limited to diminishing the protein folding burden in the ER. In mice, RIDD controls muscle regeneration by degrading mRNA of a TGFβ-family protein myostatin^46^. During development of the *Drosophila* eye, RIDD activity is required to increase the synthesis of the rhodopsin, when rhabdomeres are formed^47^. In *C. elegans* neurons, IRE1 functions as a controller of morphological change - dendritic branching during development and formation of the stress-resistant dauer larva, in XBP1-independent fashion^48–50^. Although the latter is suggestive of RIDD, the worm IRE1 has never been tested for its ability to perform the RIDD activity, and the potential targets are unknown. The paucity of molecularly characterized examples makes it hard to determine if RIDD is a commonly used mechanism in animal development or physiology.

We have examined previously published *C. elegans* transcriptomic data^51^ and identified several potential RIDD targets, including the growth factor DAF-7, a well-known developmental regulator from the TGFβ superfamily, secreted mainly from the ASI chemosensory neurons^52^. Transcription of *daf-7* is environmentally regulated, and is silenced when neurons sense unfavorable conditions - lack of food, high temperature, and high population density^52,53^. Interestingly, while secretion of DAF-7 in young larvae signals reproductive development, silencing of *daf-7* expression signals dauer transition^52,53^, a process that is genetically linked to the IRE1’s RIDD activity^48,49^. Importantly, dauers are exceptionally stress resistant, and decrease in DAF-7 even outside of the dauer signaling leads to increased resistance to heat stress and starvation and extends adult life-span^54–56^, and increases resistance to pathogens^57^.

In this study, we first use human cells to confirm that a chimeric IRE1 with the *C. elegans* cytosolic kinase/RNase domain is capable of performing the RIDD activity, and that *daf-7* mRNA is indeed a RIDD target. We then show *in vivo* that downregulation of *daf-7* mRNA by RIDD occurs upon exposure to environmental toxin tunicamycin (Tm), under conditions insufficient for detectable activation of *xbp-1* splicing. We find that downregulation of *daf-7* by RIDD provides a preemptive protective mechanism against limited food/high temperature environment, and this protection can be reversed by DAF-7 overexpression. Because Tm, like several other ER stressors, is a secreted secondary metabolite of *Streptomyces spp*., which also secrete numerous antibiotics in order to compete with other bacteria, its presence in the environment may serve as a warning for upcoming depletion of bacteria that *C. elegans* feeds on. Our data suggest that *C. elegans* sensory neurons have ‘learned’ to utilize IRE1’s RIDD activity to signal an upcoming environmental challenge and improve the population survival.

## Results

### The kinase and RNase domains of *C. elegans* IRE1 are functional in human cells

Because *C. elegans* IRE1 has not been demonstrated to have RIDD activity, we first sought to test whether *C. elegans* IRE1 can cleave a canonical RIDD target, such as the mammalian *Blos1* transcript. We chose to use a human leukemia-derived, essentially haploid cell line HAP1 that provides robust and reliable results for both RNase activities^58,59^. A subclone of this cell line with the endogenous IRE1 knocked-out via CRISPR, IRE1KO, serves as a host for testing the activity of exogenous IRE1s^58^. The kinase and RNase domains of *C. elegans* IRE1 are well-conserved relative to their human counterparts, with 79% and 83% similarity, respectively, including the key phosphorylation residues in the active loop of the kinase (marked in red, Suppl. Fig. S1A), while the lumenal and transmembrane (TM) domains are more divergent. We therefore designed a chimeric protein where the entire cytosolic portion of the human IRE1 was replaced by *C. elegans* sequences (Fig. 1A), reasoning that such configuration would avoid the potential confounding effects of the worm lumenal and TM domains needing to sense misfolded proteins and interact with the mammalian ER membranes and chaperones. The chimeric protein should thus maintain the stress recognition module of the human IRE1, while enabling functional assessment of the worm’s catalytic domains. Finally, to allow visualization of the chimeric protein in live cells, we inserted a superfolder GFP (sfGFP) between the TM and cytosolic domains (Fig. 1A), mimicking the well-characterized and fully functional human IRE1-sfGFP or -mCherry proteins^58,59^.

**Figure 1.**
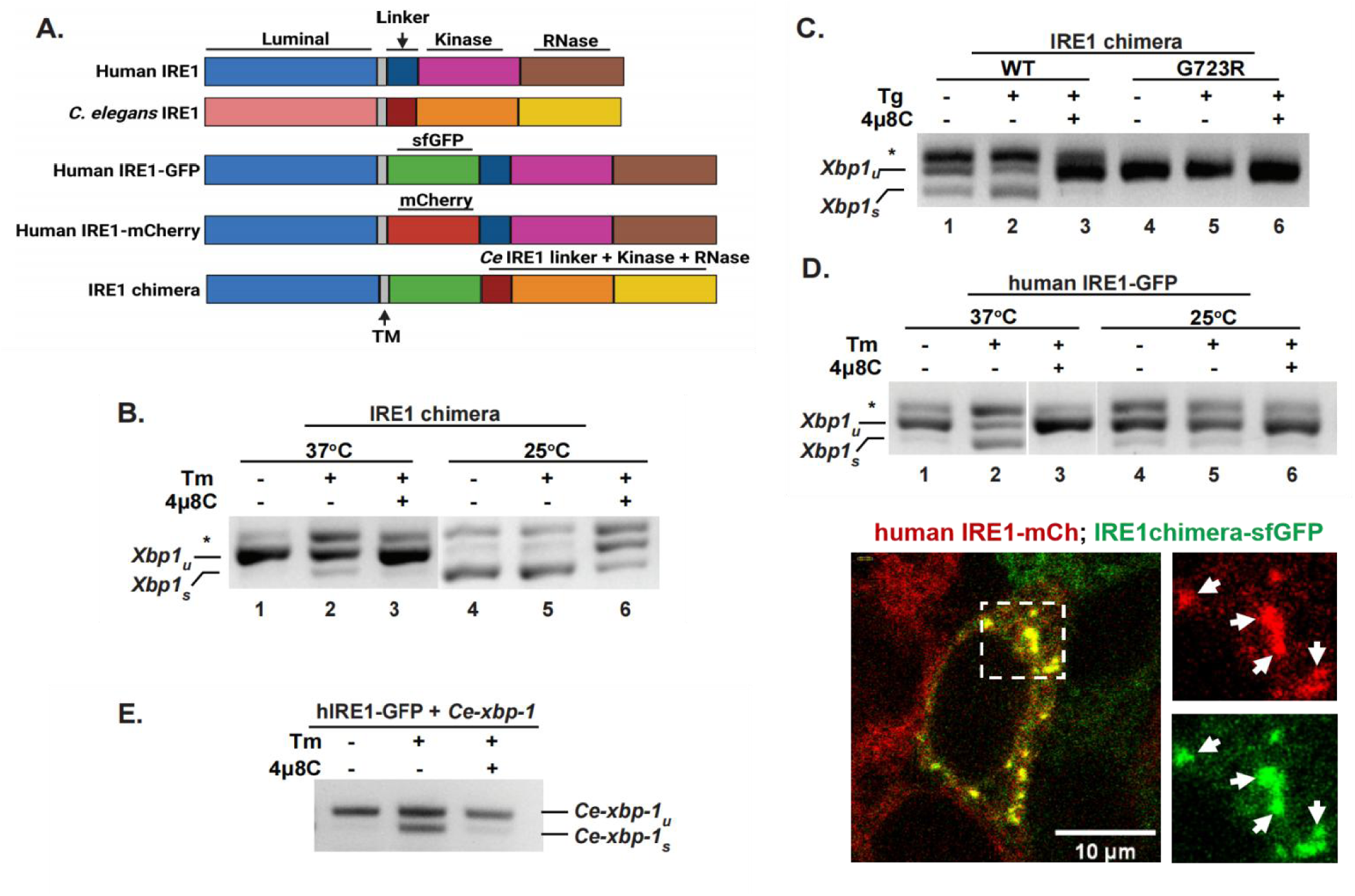
*C. elegans* cytosolic domains recognize human targets and are catalytically active in human cells. **(A)** Schemes of domain organization in the human and *C. elegans* IRE1 proteins, and the constructs used in this work. The IRE1 chimera is composed of human lumenal and transmembrane (TM) domains (amino acid 1 to 494 of human IRE1α), a super-folder GFP (sfGFP), and *C. elegans* IRE1 cytosolic portion including the linker region and the kinase/RNase domain (amino acid 456 to 967 of *C. elegans* IRE1). **(B, C)** IRE1 chimera is active and kinase-dependent. RT-PCR assay was used to measure *Xbp1* splicing in HAP1 IRE1KO cells reconstituted with an IRE1 chimera, either wildtype or G723R kinase-dead mutant. Cells were treated with 4μg/ml tunicamycin (Tm) or 2.5 μM thapsigargin (Tg) for 4hrs, either at 25°C or 37°C, as indicated. IRE1 inhibitor 4μ8C was used at 16 μM. *Xbp-1_u_:* unspliced *Xbp1; Xbp1_s_:* spliced *Xbp1;* *, non-specific PCR product, likely a hybrid of spliced and unspliced products. **(D)** RNase activity of human IRE1 is also temperature-dependent. HAP1 IRE1KO cells stably transfected with human IRE1-GFP protein were treated as in B. Human IRE1 is nearly inactive at 25°C. **(E)** Human IRE1 recognizes and cleaves *C. elegans xbp-1.* HAP1 IRE1KO cells stably expressing human IRE1-GFP protein were transfected with *C. elegans* full-length *xbp-1* and treated as in C. **(F)** IRE1 chimera is able to oligomerize and form clusters with human IRE1 under strong ER stress. HAP1 IREKO cells expressing both the human IRE1-mCherry and IRE1 chimera (containing sfGFP) were treated with SubAB (0.1μg/ml) for 4hrs. Cells were imaged at 25°C, confocal projection of a z-stack is shown for a representative cell containing clusters. Arrows: examples of clusters.

The IRE1 chimera was introduced into HAP1 IRE1KO cells through lentivirus infection and stably expressed in these cells (Fig. S1B, C). To avoid any artifacts of overexpression, we selected cell lines where chimeric proteins were expressed at levels lower than those of the fully functional^58,59^ human IRE1-GFP protein (Fig. S1C).The functionality of the chimera was first verified by RNase activity towards human *Xbp1* RNA (Fig. 1B, C). Treatment of HAP1 IRE1KO cells, expressing the IRE1 chimera, with the ER stressor tunicamycin (Tm) resulted in production of spliced *Xbp1*; the splicing was inhibited by the selective IRE1 inhibitor 4µ8C^60^, confirming it was due to the activity of the worm IRE1 domains (Fig. 1B, lanes 1-3). As the splicing activity of the chimeric protein was somewhat limited, we wondered whether worm catalytic domains were better adapted for lower temperatures, since *C. elegans* lives at the temperature range of 15°C to 25°C, lower than mammalian cells. Indeed, when HAP1 cells expressing chimeric IRE1 were treated with Tm at 25°C, a much stronger *Xbp1* splicing activity was detected, with up to 80% spliced *Xbp1* observed even without stress treatment (Fig. 1B, lanes 4-6). Importantly, this high splicing activity was still inhibited by 4µ8C. The splicing activity was also induced by a different ER stressor, thapsigargin (Tg)^61^ (Fig. 1C, lanes 1-3, Fig. S1D). Finally, to test whether the *Xbp1* splicing activity of the chimera required auto-phosphorylation, like that of mammalian IRE1, the kinase-inactivating mutation (G723R)^11^ was introduced into the chimera. The cells with mutant chimera failed to show any splicing activity even under ER stress treatment (Fig 1B). We conclude that the IRE1 chimera is functional, able to recognize and cleave a mammalian substrate, and is activated by the proper progression of kinase-RNase activation.

We asked whether the temperature-dependence of *Xbp1* splicing activity is a general property of any IRE1, by testing the activity of the human IRE1 at 25°C. We found that human IRE1-GFP essentially lost its activity and was no longer responsive to ER stress at 25°C (Fig. 1D, lanes 4-6), confirming that temperature-dependence is not a peculiar feature of the worm protein.

Finally, we asked whether cross-species recognition between IRE1 and *Xbp1* is reciprocal. We found that the human IRE1 does efficiently cleave *C. elegans xbp-1* cDNA under stress (Fig. 1E). These results indicate that human and worm IRE1s recognize and cleave their substrates in the same manner.

### Oligomerization of the IRE1 chimera and temperature dependence of IRE1

Activated yeast and mammalian IRE1 proteins exhibit a distinct manifestation of activation – clustering, reflecting their strong oligomerization^58,62^. Thus, we asked whether the chimeric IRE1 clustered in human cells, and whether it was able to co-cluster with the human protein. We found no cells with clustered IRE1 chimera under Tm stress at 25°C, when its RNAse is active (n=65). Nonetheless, IRE1 chimera could co-cluster with human IRE1-mCherry at 25°C under treatment with subAB (Fig. 1F), the most robust activator of ER stress^58^, indicating that the cytosolic domains of *C. elegans* IRE1 could adopt the conformation required to interact with human IRE1.

Interestingly, similar to the abolished splicing activity, the clustering activity of the human IRE1 protein was severely compromised at 25°C, with only ∼7% of cells (10 out of 151 cells scored) exhibiting clusters, compared to ∼85% (60 out of 71 cells) at 37°C. Strikingly, the human IRE1 clusters that were formed at 37°C dispersed within 15 mins upon shifting cells to 25°C (Fig. S2). These results suggest that IRE1 activity in both species is temperature-dependent and optimized to the growth temperature range of the host organism. Moreover, strong activation of the chimera splicing activity at 25°C without additional stressor (Fig. 1B) is consistent with a physiological UPR in worms at this temperature^63,64^, and may indicate that growth at this temperature is already sufficient to activate the enzymatic domains of *C. elegans* IRE1.

### Importance of autologous linker sequence

Considering the conservation of the enzymatic domains of the two proteins, we next wondered whether the presence of the worm linker sequences in the cytosolic portion of our chimera contributed to either the temperature sensitivity or the constitutive activity of the chimeric IRE1 protein. A linker region is located between the IRE1 transmembrane domain and kinase domain and is required for oligomerization and activation of IRE1^65^. In yeast, the linker region is also required for the recruitment and docking to IRE1 of the *Hac1* (yeast *Xbp1* homologue) substrate^66^. The human linker is considerably longer (106 aa) than the *C. elegans* one (62 aa), with no significant similarities (Fig. S1A). Notably, the mammalian linker has three Gly/Ser-rich sequences that can impart greater polypeptide flexibility, but also more Pro residues compared to *C. elegans* linker.

Thus, we exchanged the worm linker region with the human one (Fig. S1E). Strikingly, the new chimeric protein with the human linker, h-linker-IRE1 chimera, completely failed to splice *Xbp1* at either 25°C or 37°C, even under Tm stress (Fig. S1E). This indicates that an autologous linker region is required to activate the kinase and RNase domains. Thus, only the IRE1 chimera with *C. elegans* linker region was used in subsequent experiments.

### *C. elegans* IRE1 can perform RIDD

Having shown that the IRE1 chimera is functional and the cytosolic portion of *C. elegans* IRE1 can recognize and cleave human *Xbp1*, we tested if it can cleave a known RIDD substrate in HAP1 cells. mRNA of BLOC1S1 (Biogenesis of lysosome-related organelles complex 1 subunit 1) is one of the most conserved RIDD substrates, and its degradation via IRE1 RIDD activity has been shown in multiple cell lines^25,38^ upon ER stress. Therefore, we asked whether IRE1 chimera can downregulate *Blos1* mRNA. At 25°C, *Blos1* mRNA was consistently decreased by ∼25% upon Tg treatment in chimera-expressing cells, relative to mock treated cells (Fig. 2). The decreased *Blos1* reflected activation of the worm IRE1 catalytic RNase activity, since the IRE1 RNase inhibitor 4μ8C not only fully inhibited the downregulation of *Blos1* triggered by Tg treatment, but also increased the *Blos1* mRNA level by an additional ∼50%. This “upregulation” is likely due to inhibition of the spontaneous RNase activity of the chimera that we observed previously in *Xbp1* splicing assay (Fig. 1B and C). The kinase-dead mutant (G723R) chimera failed to downregulate *Blos1* mRNA (Fig. 2), further confirming the dependence on IRE1 activity. *Blos1* downregulation by ER stress was selective, since non-RIDD substrates in mammalian cells, such as *Gapdh* (Fig. S3) or mCherry (Fig. S4), were unaffected. We conclude that *C. elegans* IRE1 RNAse domain can recognize and cleave a canonical RIDD substrate.

**Figure 2.**
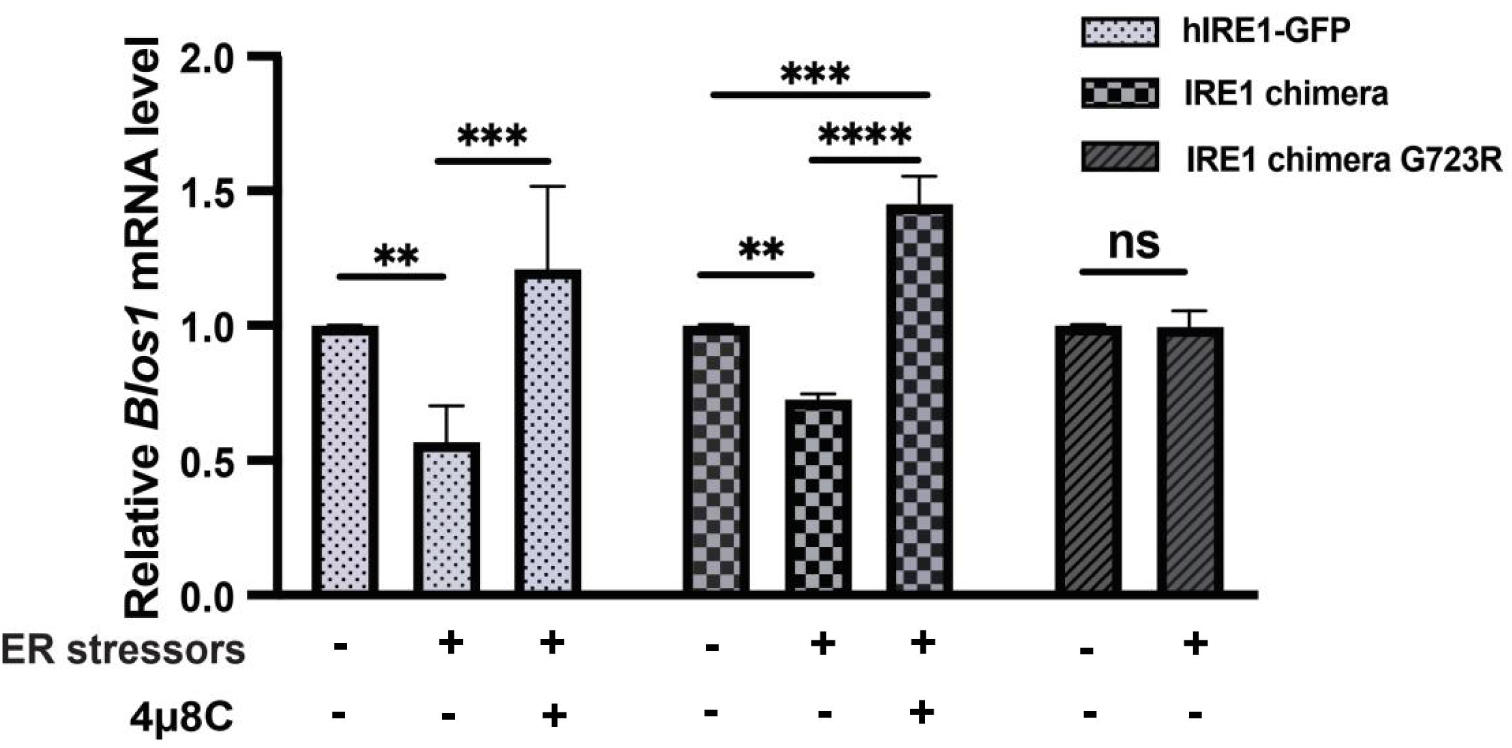
*C. elegans* IRE1 can perform RIDD activity. RT-qPCR was used to quantify mRNA levels of the canonical RIDD substrate *Blos1* in HAP1 IRE1KO cells expressing indicated IRE1 proteins. Cells with hIRE1-GFP were treated with Tm at 37°C, cells with IRE1 chimeras were treated with Tg at 25°C. *Blos1* mRNA level is normalized to the housekeeping gene *Rpl19*. 3 independent experiments, error bars: mean+/-SD, Anova with Tukey multiple comparison correction was used for hIRE1-GFP and IRE1 chimera, *t-*test was used for IRE1 chimera-G723R. Significance as described in Methods.

### *daf-7* is a RIDD substrate in human cells

With the indication that worm IRE1 is RIDD-active, we initiated a search for its endogenous RIDD substrates, based on the following expected features: their transcripts should be downregulated under ER stress, and downregulation should require the presence of IRE1 but not XBP1, and be independent of the PERK and ATF-6 branches of the UPR. We examined the previously published microarray data in Shen *et al* ^51^, using these criteria (see Methods for details). Out of 3538 transcripts with expression changes under ER stress, only 43 candidates met the above criteria (Table 1). As expected for most RIDD substrates^32,67^, 30 of the candidates were predicted to be either secretory or membrane proteins. Surprisingly, the recently identified RIDD target, *flp-6*^33^, was not among the selected candidates, as it was not downregulated upon Tm treatment in this microarray (Table 1). However, based on WormBase modENCODE data^68^, *flp-6* is mainly expressed in the early larval and dauer stages; since Shen *et al* ^51^. used non-dauer larval stage 2 (L2) animals^51^, our approach may not be sensitive enough to detect downregulation of *flp-6*. Interestingly, two of the selected candidate genes (*lect-2* and *daf-7*) are involved in dendrite arborization, which, based on the previous genetic data, may depend on RIDD^48–50^. The muscle-secreted *lect-2* contributes to the patterning of all somatosensory dendrite arbors^48,49,69,70^, while *daf-7* is linked to arborization of IL2 neurons specifically during transition into the stress-resistant dauer stage^71^. Because decrease in DAF-7 expression signals transition to the dauer stage, we focused on *daf-7* as a candidate RIDD target.

**Table 1:**
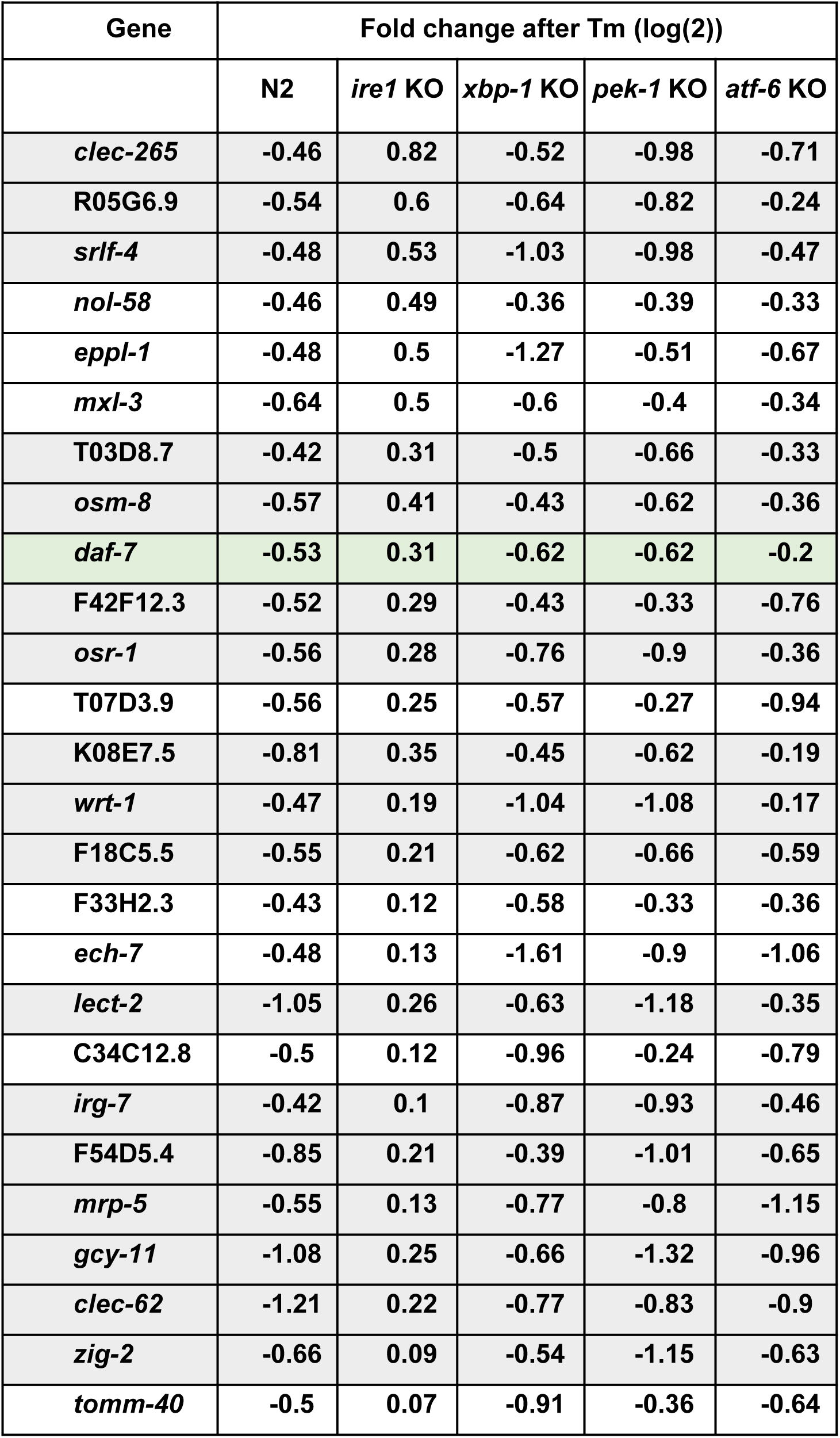
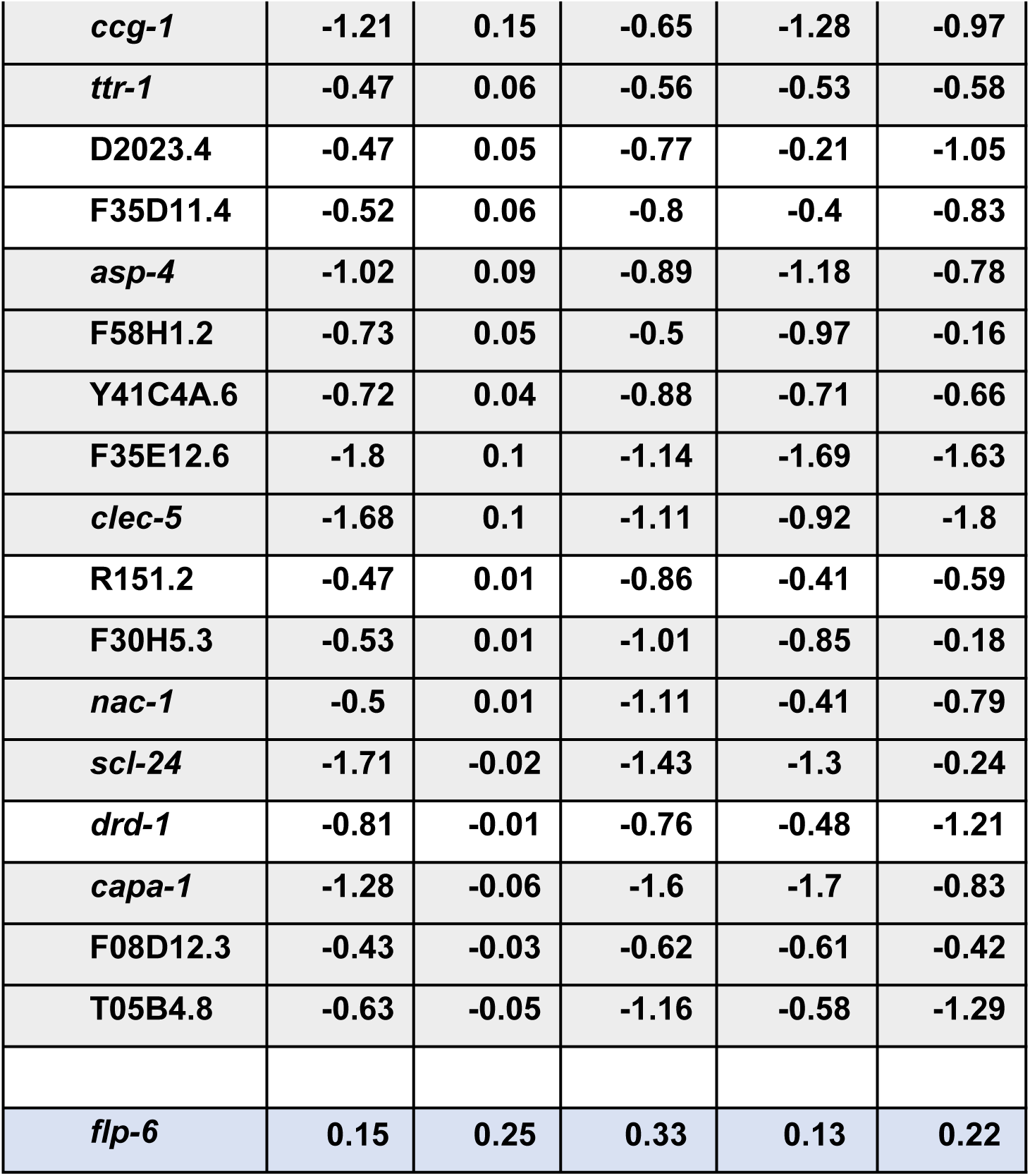
List of candidate RIDD targets, adapted from Table 5 in Shen *et al* ^51^. Transcripts are sorted by the largest difference in downregulation between wild type (N2) and IRE1-deficient animals (ΔN2-ΔIRE1KO)/ΔN2≥0.9, see Methods). Grey shading indicates transcripts coding for known secreted and membrane proteins, and those with predicted ER signal sequences. *daf-7* is shaded in green; *flp-6* (shaded blue) did not show downregulation in this microarray.

Since the above data on *daf-7* regulation by ER stress is based on genetic deletions, we sought to first independently confirm that *daf-7* mRNA can be regulated by the IRE1 RIDD activity. Thus, we compared it with the known RIDD substrate in human HAP1 cells, where the RIDD activity has been characterized^26^. Since we already showed that human IRE1 can correctly recognize and cleave the worm *xbp-1* (Fig. 1E), we expected human IRE1 to also process worm RIDD substrates, if any. We thus introduced *daf-7* mRNA into HAP1 cells, as mCherry fusion of DAF-7 protein (Fig. 3A) that we have previously demonstrated in *C. elegans* to be fully functional^72^.

**Figure 3.**
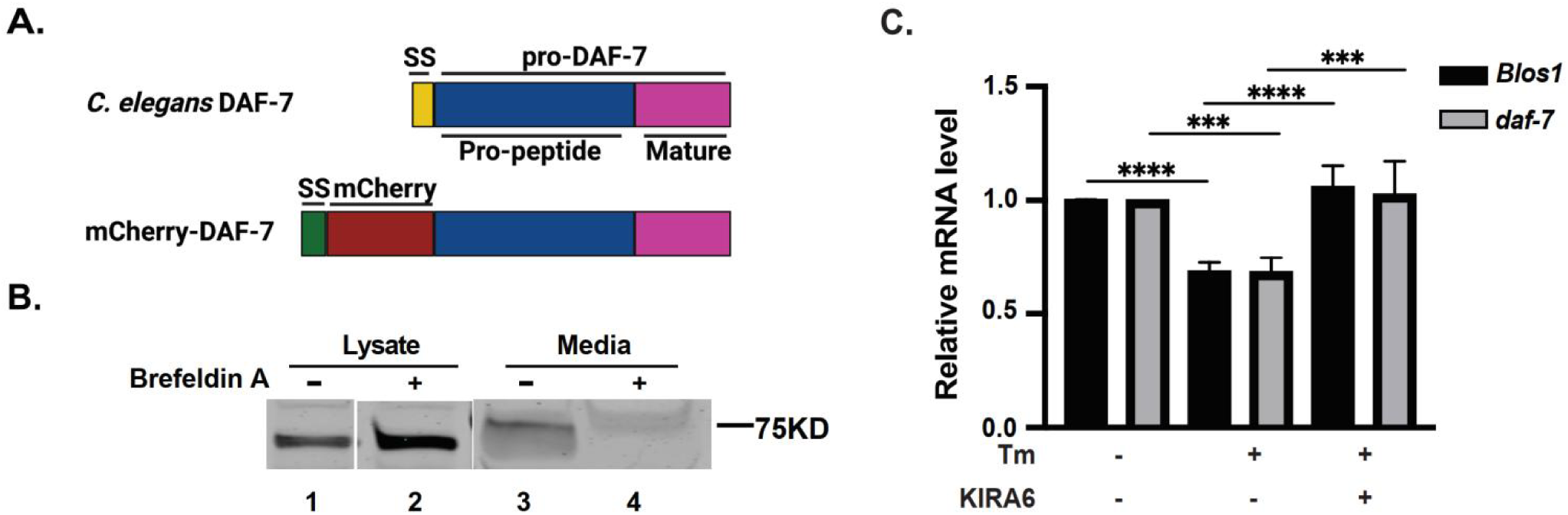
*daf-7* is degraded by the human IRE1 as a RIDD substrate. **(A)** Scheme of the domain composition of DAF-7 protein and the construct used in this work. *C. elegans* DAF-7 contains a signal peptide (SS) and pro-DAF-7 (aa. 23 to 350 of DAF-7), including latency-associate peptide (Pro-peptide) and mature DAF-7. The mCherry-DAF-7 construct contains human IGF2 signal peptide (aa. 1 to 24 of IGF2) in place of the original DAF-7 signal peptide, followed by mCherry and pro-DAF-7 (aa. 23 to 350 of DAF-7 protein), and a C-terminal V5/6-His tag. **(B)** mCherry-DAF-7 is secreted in HAP1 cells. mCherry-DAF-7 was stably expressed in HAP1 WT cells, cells were treated with Brefeldin A (2μg/ml) for 24hrs to block secretion, and analyzed by Western blot with anti-mCherry antibody. (C) *daf-7* mRNA is degraded by human IRE1 under ER stress. RT-qPCR analysis of *Blos1* and *daf-7* mRNA levels in HAP1 cells expressing mCherry-DAF-7. Cells were treated with Tm (4μg/ml) and IRE1-specific inhibitor KIRA6 (1μg/ml) for 6hrs, 3 independent experiments, error bars: mean+/-SD, Anova with Tukey multiple comparison correction, significance as described in Methods.

To ensure that *daf-7* mRNA can be recognized by translation elements and targeted to the ER, we replaced DAF-7 signal peptide with that of human IGFII. ER targeting was confirmed by secretion to the culture media, which was sensitive to Brefeldin A (Fig. 3B). We then treated HAP1 cells expressing mCherry::DAF-7 with Tm, and assayed *daf-7* mRNA levels. *daf-7* message was downregulated to the same extent as the canonical RIDD substrate *Blos1* (31+/-2% and 32+/-3% respectively, n=4) (Fig. 3C). Furthermore, the downregulation of both *daf-7* and *Blos1* was suppressed by IRE1 inhibitors, KIRA6, a type II kinase/IRE1inhibitor^73^ (Fig. 3C), and 4μ8C. Importantly, the downregulation reflected cleavage of *daf-7* sequences within the mCherry::DAF-7-coding transcript, since *mCherry* sequence was unaffected by either Tm or IRE1 inhibitors treatments (Fig. S4). Thus, *daf-7* mRNA is degraded by the human IRE1 like a RIDD substrate.

As mentioned above^22,25,32,38–40^, RNAs degraded by RIDD activity may or may not have a *Xbp1-*like spatially conserved CNG↓CNGN IRE1 recognition sequence^41^ *C. elegans xbp-1* message contains two such loops that are specifically cleaved by IRE1 during splicing^11^ (Fig. S5A). The RNA fold prediction algorithm^74^ identified a stem-loop in *daf-7* mRNA with a potential IRE1 cleavage site (Fig. S5A, right) . The size of the loop is 9nts, larger than the typical 7nts. We asked whether this was the IRE1 recognition site, by mutating conserved residues on both sides of the predicted cleavage site. Incidentally, one of the mutations also eliminates a cysteine residue in the growth factor domain (C314S). Tm treatment still downregulated both mutated *daf-7* mRNAs with similar efficiency to the *Blos1* (Fig. S5B, C), suggesting either the existence of more than one cleavage sites, or of a different type of a cleavage site(s).

### *daf-7* is downregulated by IRE1 *in vivo* at correct developmental stages and under physiological stress levels

The data so far show that *C. elegans* IRE1 is capable of performing a RIDD activity (Fig. 2), that

*daf-7* message is downregulated by Tm stress in mixed populations of *C. elegans* in a manner consistent with it being a RIDD target^51^ (Table 1), and that *daf-7* mRNA is degraded by a human IRE1 as a RIDD substrate (Fig. 3). To understand the possible physiological significance of *daf-7* regulation by RIDD, we considered that decrease in DAF-7 secretion signals the presence of stressful environment specifically in late L1 (first larval stage)/early L2 animals. Thus, we tested whether activation of IRE1 can reduce *daf-7* transcript levels at these stages. When synchronized late L1 populations were treated for 2 hours with Tm under conditions that activate robust IRE1-dependent *xbp-1* splicing^75^, the transcript levels of *daf-7* were strongly downregulated by 48% +/-29% (Fig. 4A, n=6). This was not due to non-specific decrease in transcription, since two unrelated housekeeping genes, *ama-1* and *acs-20* (Fig. 4A), as well as the transcript of another neuro-secretory protein, *ins-1* (Fig. S6A), were not affected. As expected for IRE1-dependent process^75^, the downregulation of *daf-7* begun to attenuate at 4hrs Tm treatment (Fig. 4A), paralleling the *xbp-1* splicing kinetics in the same animals (Fig. 4B, C). Importantly, the downregulation of *daf-7* was dependent on IRE1 activity, as Tm treatment of larvae carrying enzymatically inactive mutant of IRE1 (G723R, *zc14* allele)^11^ failed to downregulate *daf-7* transcript, and in fact increased its levels (Fig. 4A). Conversely, the downregulation of *daf-7* was still observed in larvae without functional XBP-1 (Fig. S6B). This indicates that *daf-7* message is a selective RIDD target of activated IRE1 at the developmental stage when it functions to signal stressful environment.

**Figure 4.**
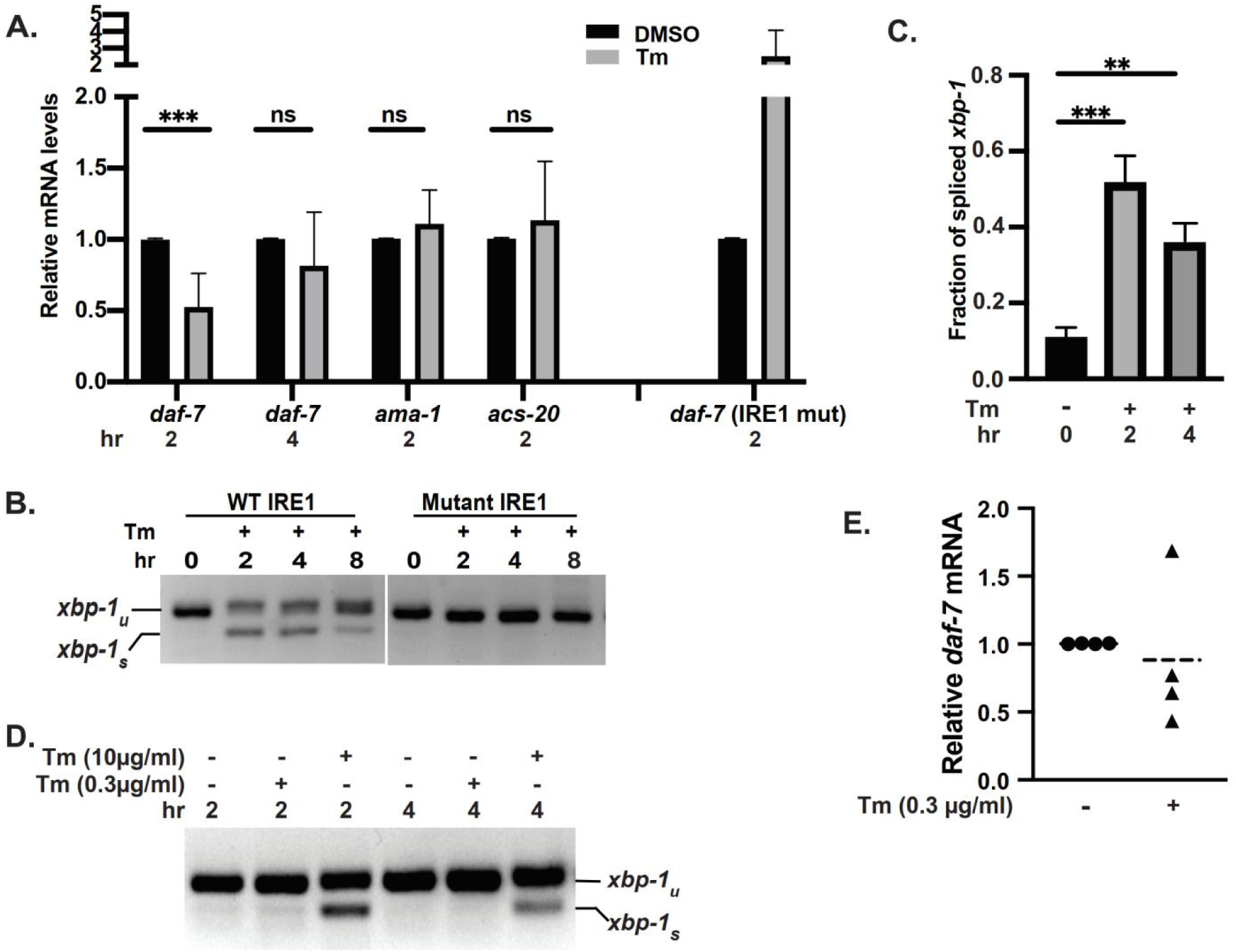
*daf-7* mRNA is a RIDD substrate *in vivo*. **(A)** *daf-7* mRNA is selectively downregulated by IRE1 in *C. elegans* under ER stress. RT-qPCR analysis of *daf-7* mRNA levels in synchronized late L1 larvae, normalized to actin. Animals were treated with Tm (10μg/ml) for 2hrs or 4hrs. Mutant IRE1: SJ30(*zc14*) animals carrying kinase-dead IRE1(G723R). *daf-7:* 6 (2hrs) or 4 (4hrs) independent experiments; *acs-20* and *ama-1*: 3 experiments. IRE1 mut: 4 experiments. Error bars: means +/-SD, *t-*test, significance as described in Methods. **(B, C)** Time-course of UPR induction. RT-PCR *xbp-1* splicing assay in late L1 larvae treated with Tm (10μg/ml) for indicated number of hours. **C:** quantitation in animals with wild type IRE1. 3 independent experiments, error bars: mean+/-SD, Anova with Tukey multiple comparison correction. **(D)** Marginal activation of *xbp-1* splicing at low Tm concentration. Late L1 larvae were treated with high (10μg/ml) or low (0.3μg/ml) Tm concentration for 2 or 4hrs. Representative RT-PCR from 3 (high conc.) and 4 (low conc.) experiments. **(E)** *daf-7* mRNA is still downregulated at low Tm concentration. RT-qPCR analysis of *daf-7* mRNA levels in late L1 larvae treated with 0.3μg/ml Tm for 4hrs. 4 independent experiments shown.

We next considered whether degradation of *daf-7* message was only triggered under severe ER stress. We treated synchronized late L1 animals with 0.3μg/ml Tm, a 100-fold lower concentration than was used previously to induce ER stress in worms^51^. Treatment with 0.3μg/ml Tm resulted in *xbp-1* splicing at only barely detectable levels at 2 hours, and no detectable splicing at 4 hours (Fig. 4D). Strikingly, even at this low Tm concentration, *daf-7* mRNA was downregulated in 3 out of 4 independent experiments (Fig. 4E), to the levels comparable to those triggered by high Tm (Fig. 4A). Downregulation of *daf-7* mRNA by the low concentration of Tm was selective, since non-RIDD target *ins-1* was unaffected (Fig. S6A). Thus, RIDD of *daf-7* is the preferred IRE1 activity under low ER stress.

### Downregulation of *daf-7* message by Tm is not sufficient to induce dauer transition

The best-defined phenotype expected from reducing DAF-7 levels is dauer larvae formation. Thus, we tested dauer induction by combining a standard treatment of crowding-induced starvation and higher temperature (25°C)^76^ with an added dose of Tm. In mock-treated (DMSO) wild type animals, dauer larvae started to appear within 48hrs after starvation and their number increased by 96 hrs (Fig. S7, left). Addition of Tm not only failed to increase the number of dauer larvae, but inhibited dauer formation, and this was true even at very low Tm concentration of 0.1μg/ml (Fig. S7). This suggests that downregulating *daf-7* by RIDD does not alone induce dauer larvae. Since even at high concentration (10μg/ml), Tm only downregulated *daf-7* mRNA by ∼50% (Fig. 4A), RIDD activity may not reduce *daf-7* levels enough to trigger dauer transition. Alternatively, Tm may prevent dauer transition by interfering with various required secretory events^77,78^.

### Downregulation of *daf-7* by RIDD allows for survival and growth of *C. elegans* populations in limited food/high temperature environments

While performing dauer experiments, we noticed that worm populations under higher temperature grew better when treated with low Tm concentrations compared to non-Tm controls. Thus, we considered environmental conditions under which the animals could encounter Tm. Although Tm is commonly used to experimentally activate the UPR in eukaryotic cells, it is a natural secondary metabolite antibiotic secreted by several *Streptomyces spp*.^79^. Moreover, a number of other compounds that are used to trigger stress in eukaryotic cells, including such common UPR inducers as lactacystin, ionomycin, and calcium ionophore, are in fact antimicrobials secreted by *Streptomyces*. The presence of *Streptomyces* may thus negatively impact worm populations not only by secreting nematocidal metabolites, but by depleting other bacteria that serves as food source for *C. elegans*. Moreover, reproduction of *C. elegans*, and thus population growth, is inhibited at temperatures that are appropriate for *Streptomyces* (27°C or higher)^80,81^ (Fig. 5C). Because deletion of *daf-7* is known to increase larval survival of starvation and high temperature streses^55^, we asked if downregulation of *daf-7* message by RIDD could be protective under conditions mimicking the presence of *Streptomyces* - a combination of 27°C and sparse food availability. Embryos were placed sparsely on Tm-containing plates and allowed to grow until plates were crowded with next generation larvae and food became limiting. Strikingly, inclusion of low concentrations of Tm (0.1 and 0.3μg/ml) rescued the population growth defect under these conditions, increasing the population size (Fig. 5A) and the number of laid embryos (Fig. S8A) by 2-3 folds within 72 hours. This rescue continued into the next generation of animals, despite complete depletion of food, since low concentrations of Tm still increased the population size more than two-fold after 96 hours (Fig. S8B).

**Figure 5.**
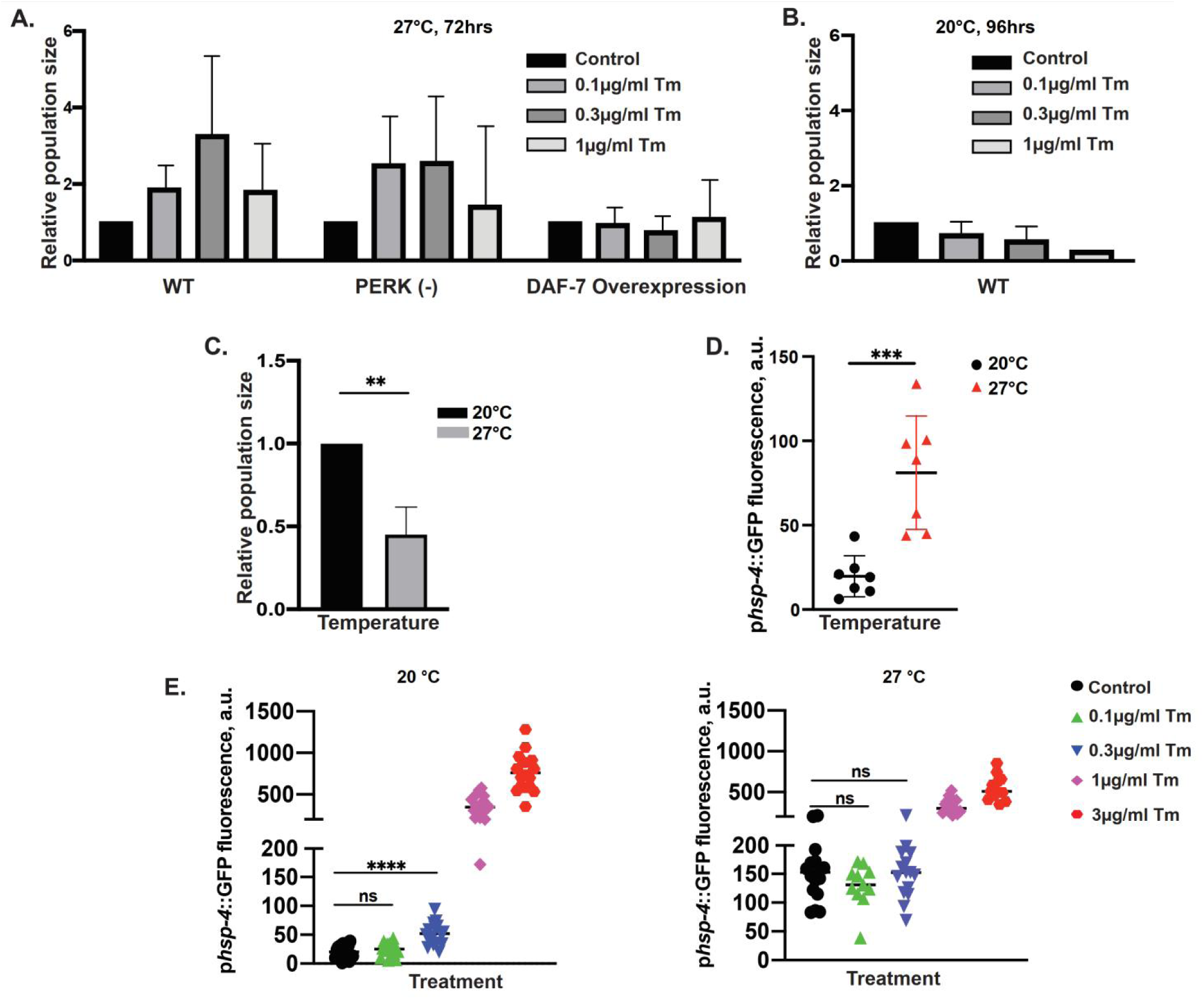
Pre-emptive downregulation of *daf-7* by RIDD supports survival of nutritional/temperature stress. **(A)** Pretreatment with low Tm concentrations increases population growth under stressful environment through downregulation of DAF-7. Population size assays: animals of indicated genotypes were exposed to Tm or DMSO (control) starting from embryo stage, under low population density (25 embryos per 3cm plate) and plentiful food. Animals were allowed to grow until plates were crowded and food nearly depleted (72hrs) at 27°C. PERK(-): *pek-1(ok275)*; DAF-7 overexpression: animals transgenic for *pdaf-7::mcherry::daf-7.* 3 independent experiments, error bars: mean+/-SD. **(B)** Low Tm is not beneficial at normal growth temperature. Wild type embryos were plated as in A and allowed to grow until plates were crowded and food nearly depleted (96hrs) at 20°C. 3 independent experiments, error bars: mean+/-SD. **(C)** Elevated temperature (27°C) restricts population growth. Embryos were plated as in A and population sizes measured after 72hrs growth at 27°C or 96hrs at 20°C for. 3 independent experiments, error bars: mean+/-SD, *t-*test, significance as described in Methods. **(D)** Induction of UPR reporter by growth at elevated temperature (27°C). GFP fluorescence in late L1 animals carrying p*hsp4::*GFP reporter, grown at 20°C or 27°C from embryo stage. error bars: mean+/-SD, *t-*test. **(E)** Elevated temperature (27°C) has a dominant effect on UPR reporter induction at low Tm concentrations. Animals were treated (as in D). Comparison of control (DMSO) and low concentration Tm (0.1µg/ml and 0.3µg/ml) treatments: Anova with Tukey multiple comparison correction.

Importantly, overexpression of DAF-7 completely abolished the rescue effect of Tm, indicating that decrease in *daf-7* was required for the rescue (Figs. 6A and S9). Notably, the rescue was specific to the growth in these otherwise restrictive conditions, since even low concentrations of Tm were detrimental at the normal growth temperature of 20°C (Fig. 5B). Thus, Tm- and RIDD-mediated decrease in *daf-7* can serve as an adaptation strategy to ensure population survival in stressful environments.

### Hormesis or PERK activity does not account for the rescue of survival by low Tm

Unfortunately, we were unable to directly test whether the rescue of population growth by Tm exposure requires IRE1 or XBP1, since animals deficient for either of these proteins arrest early in development at elevated temperatures (ref^63^ and our observations). Thus, we examined the possible alternative explanations. First, low Tm concentrations at elevated temperature could induce higher levels of general ER stress, in which case the rescue could be due to a hormesis-like effect of UPR^82,83^, rather than depending on decrease of DAF-7. We measured the fluorescence of the well-characterized p*hsp-4*::GFP UPR reporter^11^ and found that the temperature alone was sufficient to increase the reporter fluorescence (Fig. 5D), in agreement with the chimeric IRE1 activation in human cells (Fig. 1B,C) and physiological UPR in worms^64,84^. However, since the temperature effect appeared to be dominant, and the rescuing Tm concentrations (0.1 and 0.3μg/ml) did not increase the reporter fluorescence compared to the DMSO control at 27°C (Fig. 5E), these data are inconsistent with hormesis effect.

Second, we tested the possibility that the related stress sensor, PERK, mediated the rescue by low Tm. Previously, induction of strong ER stress in the ASI neuron in combination with elevated temperature was shown to trigger dauer transition^85^, which may provide survival advantage under stress. The dauer induction was shown to be due to translational silencing of the ASI neuron *via* PERK-mediated eIF2α phosphorylation; since the ASI neuron is the main source of DAF-7 for dauer signaling, activation of PERK by Tm could be responsible for our rescue observation. However, when PERK-deficient worms (*ok275*) were subjected to the same regime as the as the wild type animals, low Tm dose still increased the size of the population at 27°C by more than 2-fold (Fig. 5A). Thus, rescue of population growth by low Tm treatment is independent of PERK. Combined with the inability of Tm to rescue animals that over-express DAF-7, these data argue that activation of RIDD activity of IRE1 in sensory neurons by environmental stress promotes survival of the worm population via degradation of *daf-7* message.

## Discussion

The major conclusions from this study are that in addition to the *xbp-1* splicing activity, *C. elegans* IRE1 has a RIDD activity that is similar to that of mammalian IRE1; that *daf-7* is a physiological RIDD target; and that downregulation of *daf-7* levels by the RIDD activity modulates neuroendocrine output of sensory neurons and thus allows for survival of worm populations under conditions mimicking the presence of *Streptomyces spp.* capable of killing the bacteria that worms feed on^86^. This protection is achieved not through the already known mechanisms such as translational silencing of neurons^85^, XBP-1-dependent neuronal signaling^87,88^, or hormesis^83,89^. Previous genetic data implicated IRE1 RIDD activity in morphological remodeling of neurons in context of dauer transition^49^. We propose that pre-emptive activation of RIDD in environmentally-exposed sensory neurons of *C. elegans* acts as signaling mechanism for protection against adverse environmental conditions, including anticipated nutrient depletion and temperature stress.

### *C. elegans* IRE1 is capable of RIDD

Since IRE1 RIDD activity is not well-defined in *C. elegans*, and there is only one verified RIDD target, *flp-6*^33^, we tested *C. elegans* IRE1 RIDD using a human/*C. elegans* chimera in mammalian cells. We show that *C. elegans* RNase domain can process the heterologous RIDD substrate *Blos1*^38^, with the same efficiency and specificity as that of the human IRE1. *Blos1* often serves as a diagnostic RIDD substrate since, unlike many other mammalian RIDD targets, it is universally degraded by the activated IRE1 in different cells. Similarly, both human and *C. elegans* IRE1 efficiently and specifically degraded the newly-identified worm RIDD target, *daf-7.* RIDD of *daf-7* can operate *in vivo* during the time of developmental transition between the larval L1 to L2 stage, which is orchestrated by secretion of DAF-7 protein^52^, suggesting a physiological role for the worm RIDD activity. During this time, *daf-7* exhibits a sharp peak of expression^68^, making its levels sensitive to tuning either by the well-characterized transcriptional regulation^52,53^, the poorly understood control of its release, or the newly discovered here regulatory modality - degradation of its mRNA by the RIDD activity.

A strong argument for the physiological relevance of the RIDD activity of *C. elegans* IRE1 is that it is activated in the environmentally-exposed sensory neurons under conditions of low tunicamycin exposure, prior to the measurable induction of *xbp-1* splicing. This is important, because high doses of this toxin cause massive ER stress, which can force artificial enzyme activation. Moreover, tunicamycin inhibits glycosylation in eukaryotic cells, which can in itself interfere with normal signaling by various growth factors^90^. We observed a protective effect of pre-emptive exposure to 100-300 times lower amounts of tunicamycin that those typically used to induce ER stress. Intriguingly, both the stressful environment by itself and the low tunicamycin in the absence of stressful environment are inhibitory for the growth and survival of the population, while pre-exposure to tunicamycin decreases these inhibitory effects. Thus, the neurons may have ‘subverted’ this cellular homeostatic stress response as stress-signaling modality.

### Environmental and structural requirements

In addition to the cross-recognition of each other’s RIDD substrates, the *C. elegans* IRE1 and associated machinery processed human *Xbp1* and conversely, *C. elegans xbp-1* was processed by the human machinery. This cross-species recognition has also been observed *in vitro* between mammalian and yeast^91^ and between human and *C. elegans*^33^ IRE1 and substrates. On the other hand, we observed a complete dependence of enzyme activity on the correspondence between the linker and kinase/RNase domains: the chimera was only functional when the linker was cognate to the catalytic domains, rather than to the rest of the molecule (TM and lumenal domains) or the host cell. The reason for this is unclear at present. The linker region could transfer the conformational change from the lumenal domain, or contain yet unknown functionally important regulatory sites. The different length or amino acid composition may impart distinct biophysical properties to the linkers, which may then affect the formation of a face-to-face kinase interface for dimerization and auto-phosphorylation^65^, or a back-to-back interface for oligomerization^4^.

IRE1 proteins from two species exhibited sharp temperature dependence - the chimera had more robust RNase activity at 25°C compared to 37°C, while the human IRE1 in the same experiment was active at 37°C and inactive at 25°C. Although these data are limited to two species, it is tempting to speculate that IRE1 activation is adapted to the animal’s environment, as is also suggested by the induction of worm’s canonical UPR gene, the inducible BiP/*hsp-4*, by high growth temperatures^64,84^. This IRE1 adaptation is different from the classical model of enzyme thermo-sensitivity, in which elevated temperatures increase the rate of reaction until denaturing temperature is reached. Indeed, *C. elegans* enzymatic domains have much lower optimal temperature, yet remain activatable at the high temperatures. It is more compatible with recently proposed three-state model, where an active and inactive (but activatable) states of an enzyme are in reversible equilibrium^92^; the sequence variation between species may determine the parameters of such equilibrium. The rapid switching between active and inactive forms that we observed in the temperature shift experiments support the idea of reversible equilibrium.

### *daf-7* is a physiological RIDD target

After myostatin, DAF-7 is now a second member of the TGFβ family to be physiologically regulated by RIDD^46^. To identify candidates in *C. elegans*, we took advantage of previously performed transcriptional analysis of UPR in strains deficient for individual sensors/effectors^51^. The candidates conform to expectations of a limited number of strong RIDD substrates (43 out of 3538 transcripts in the array), and of enrichment for ER-targeted transcripts of secretory proteins (∼70%). Since two of the strong candidates, *daf-7* and *lect-2*, participate in the morphological remodeling of dendrites during transition into dauer, and we show that RIDD tunes *daf-7*’s signaling at developmental stage when the dauer decision is made, RIDD activity of IRE1 may have an unappreciated role in regulating the developmental and behavioural decisions at this crucial developmental point.

Downregulation of *daf-7* mRNA by RIDD activity is partial, and thus it regulates DAF-7 signaling differently than a complete loss of function or gene deletion. Different degrees of DAF-7 loss lead to distinct phenotypes^55,93^. The worms without both *daf-7* alleles enter only the pre-dauer L2d state if grown at 20°C with plentiful food, while they enter dauer at almost complete penetrance at 25°C^52,72^; heterozygous animals with one wildtype *daf-7* allele show no dauer phenotype even at 25°C. Importantly, DAF-7 deletion mutants exhibit increased lifespan and increased resistance to heat stress; this lifespan extension is not observed in animals treated with *daf-7* RNAi, which partially downregulate DAF-7^55^. Therefore, physiological roles of DAF-7 signaling are protein dose-dependent. While the partial loss of DAF-7 due to RIDD is insufficient to initiate dauer transition, we find that it causes a novel, paradoxical phenotype, where treatment with a toxin results in improved animal survival under restrictive high temperature (27°C).

### A physiological function of *daf-7* RIDD: signaling impending stress

RIDD can promote survival of cells enduring ER stress, either by inhibiting apoptosis^23^, or by improving stress resistance^29^. Furthermore, RIDD can be protective even before initiation of UPR, by reducing levels of transcripts of high flux proteins to prevent ER overload. Such anticipatory response is important for differentiation of secretory cells, like the insulin producing or antibody-secreting cells^94,95^. We argue that RIDD of *daf-7* fits into the protective role by preparing *C. elegans* for the impending temperature/nutritional stresses when encountering predatory bacteria, such as *Strepomyces spp.*^86^. These predatory bacteria threaten the food supply of the nematode by secreting antibiotics that kill other bacteria that worms feed on^96^. We note that secreted antibiotics have been suggested to act as signaling molecules in microbial communities^97^. As some of these antibiotics, like tunicamycin, lactacystine and ionomycin^96,98^, are ER stressors^99–102^, *C. elegans* can exploit them as warning signals for anticipated starvation.

The presence of such ‘warning signals’ appears to activate IRE1 RIDD and downregulate *daf-7* mRNA even at low concentrations that, by themselves, are not sufficient to trigger a global ER stress response. We propose that this preferential activation of IRE1’s RIDD activity in chemosensory neurons that are exposed to the nematode environment serves a signaling function that coordinates protection from upcoming stress. A similar anticipatory protection has been described for neuronal sensing of temperature, which allows induction of the heat-shock response in non-neuronal tissues prior to protein misfolding^103^, as well as for olfactory detection of pathogenic bacteria, which triggers UPR^104^ and primes HSF-1^105^. Worm neurons can detect *Streptomyces* and trigger avoidance behaviour^106^; the RIDD activity in sensory neurons modulates their neuroendocrine signaling to coordinate pro-survival strategy.

## Acknowledgments

This work was supported by the NIH grant AG063029 to TG and YA. Some strains were provided by the Caenorhabditis Genetics Center, which is funded by NIH Office of Research Infrastructure Programs (P40 OD010440).

## Supplementary Figures

**Figure S1.**
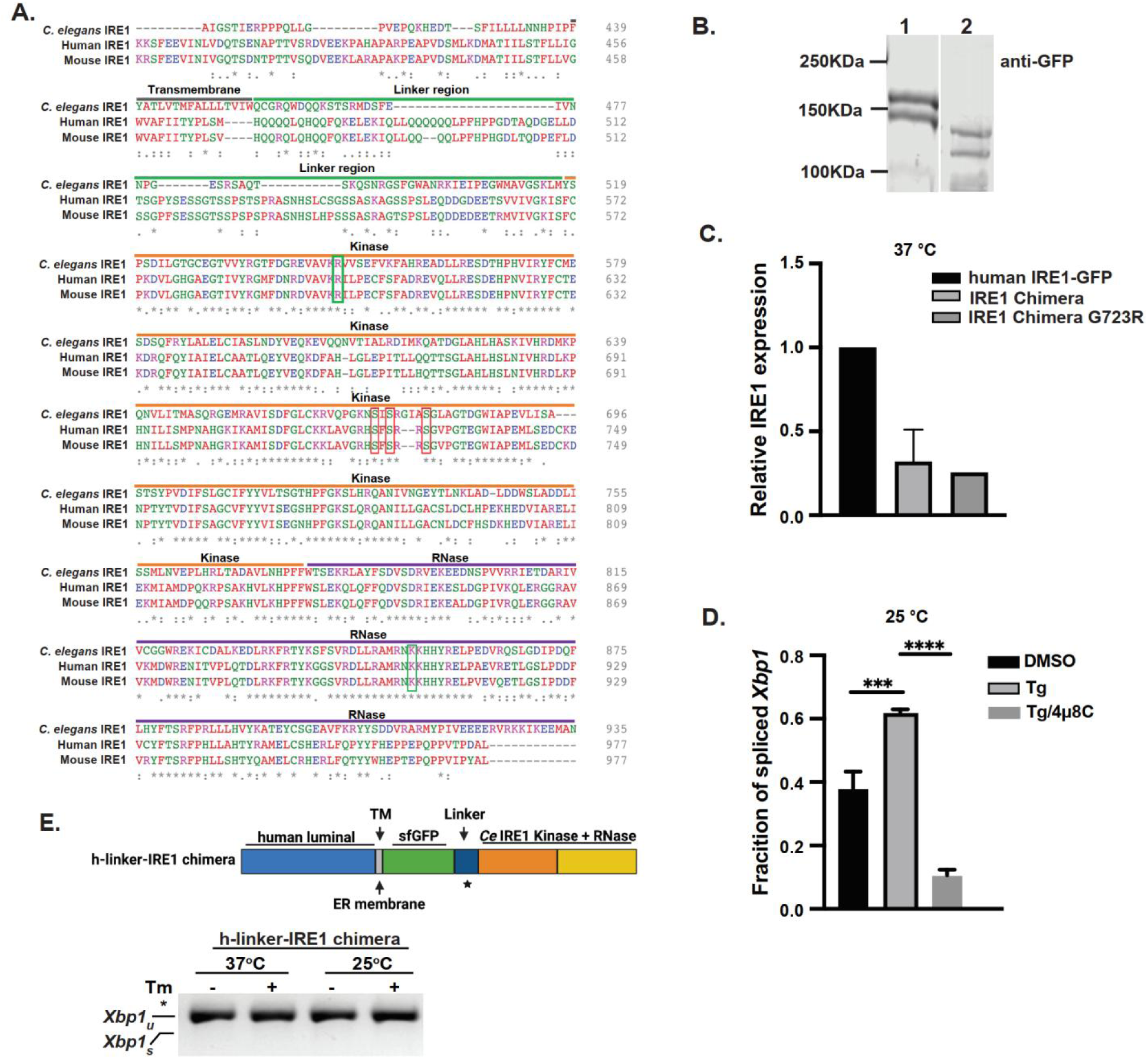
IRE1 chimera with *C. elegans* linker region is functional in human cells. Related to Figure 1. **(A)** Alignment of cytosolic domains of mammalian and *C. elegans* IRE1s (Clustal Omega^107^). **\***, indicates fully conserved; :, highly similar; **·**, weakly similar. Partial transmembrane domain (grey), linker region (green), kinase domain (orange) and RNase domain (purple) of *C. elegans* IRE1 are indicated. Red boxes: serine residues in the activation loop of kinase domain; green boxes: 4μ8C contact sites. **(B and C)** Western blot (with anti-GFP) and quantitation (with anti-N-term IRE1) of transgenic IRE1 proteins stably expressed in HAP1 IRE1KO cells. Tubulin was used as a loading control. **(D)** IRE1 chimera is active in *Xbp1* splicing in human cells, and its basal activity is augmented by Tm and inhibited by IRE1-selective inhibitor 4μ8C. Cells were treated with Tm (4μg/ml) for 4hrs at 25°C; splicing activity was analyzed by RT-PCR assay.Anove with Tukey correction, n=3. **(F)** IRE1 chimera with human linker is not functional. Protein with human IRE1 linker region (star) was expressed in HAP1 IRE1KO cells. Cells were treated and analyzed at indicated temperatures, as in D.

**Figure S2.**
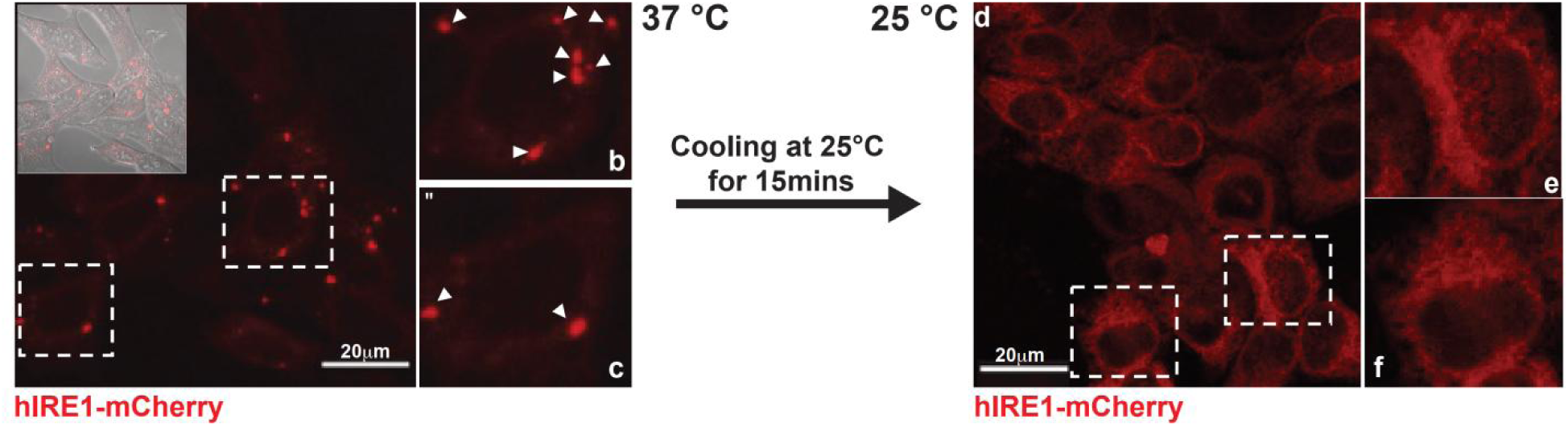
IRE1 clusters are rapidly dispersed upon temperature downshift. Related to Figure 1. HAP-1 IRE1KO cells expressing human IRE1-mCherry were treated with Tm (4μg/ml) at 37°C and then cooled for 15 minutes at 25°C . Maximum intensity projections of confocal z-stacks are shown; scale bars = 20μm. Arrowheads point to clusters; inset in 37°C image shows DIC image; close-ups are of individual cells.

**Figure S3.**
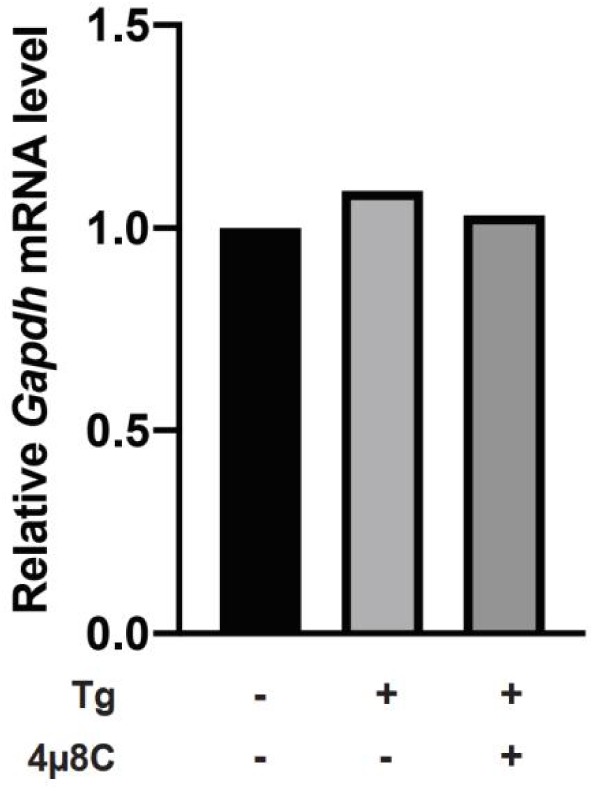
IRE1 activation does not cause general downregulation of mRNA. Related to Figure 2. RT-qPCR analysis of non-RIDD substrate *Gapdh* mRNA levels in the HAP1 IRE1KO cells expressing IRE1 chimeras. Cells were treated with 2.5μM Tg and 16μM 4μ8C for 4hrs at 25°C. *Gapdh* mRNA was normalized to *Rpl19*. Means of 2 independent experiments.

**Figure S4.**
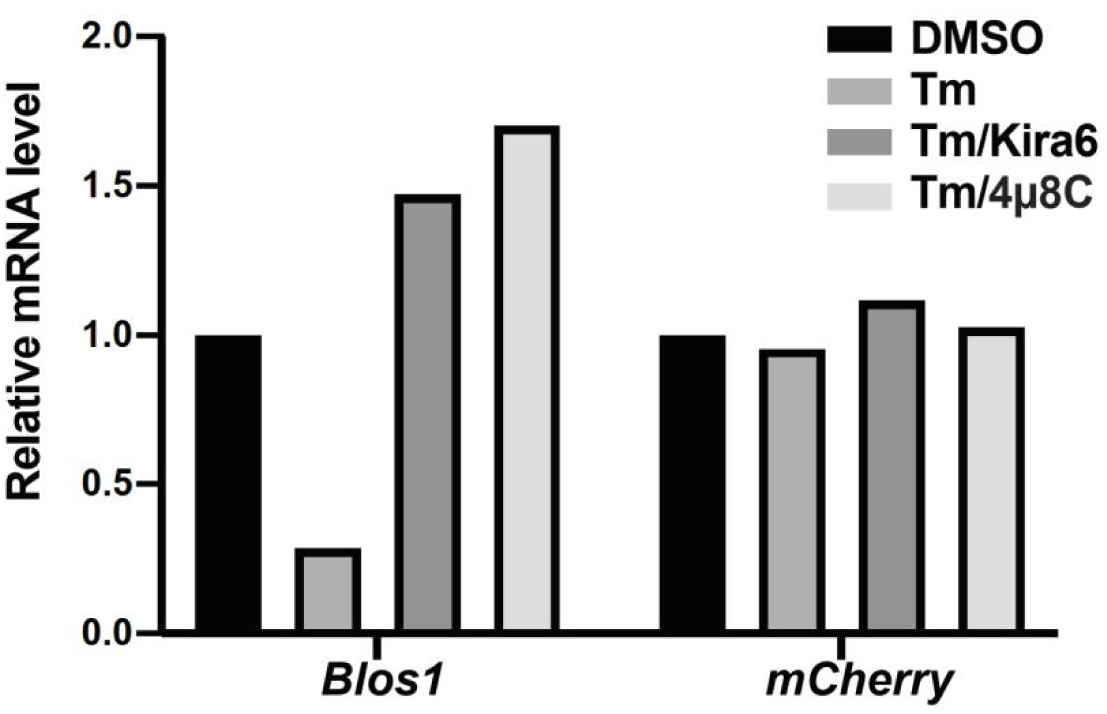
*mCherry* is not a RIDD target. Related to Figure 3. RT-qPCR analysis in HAP1 IRE1KO cells expressing human IRE1-GFP. Cells were treated with Tm (4μg/ml) and IRE1-specific inhibitors KIRA6^73^ (1μg/ml) and 4µ8C (16μM) for 6hrs. Normalized to *Rpl19*

**Figure S5.**
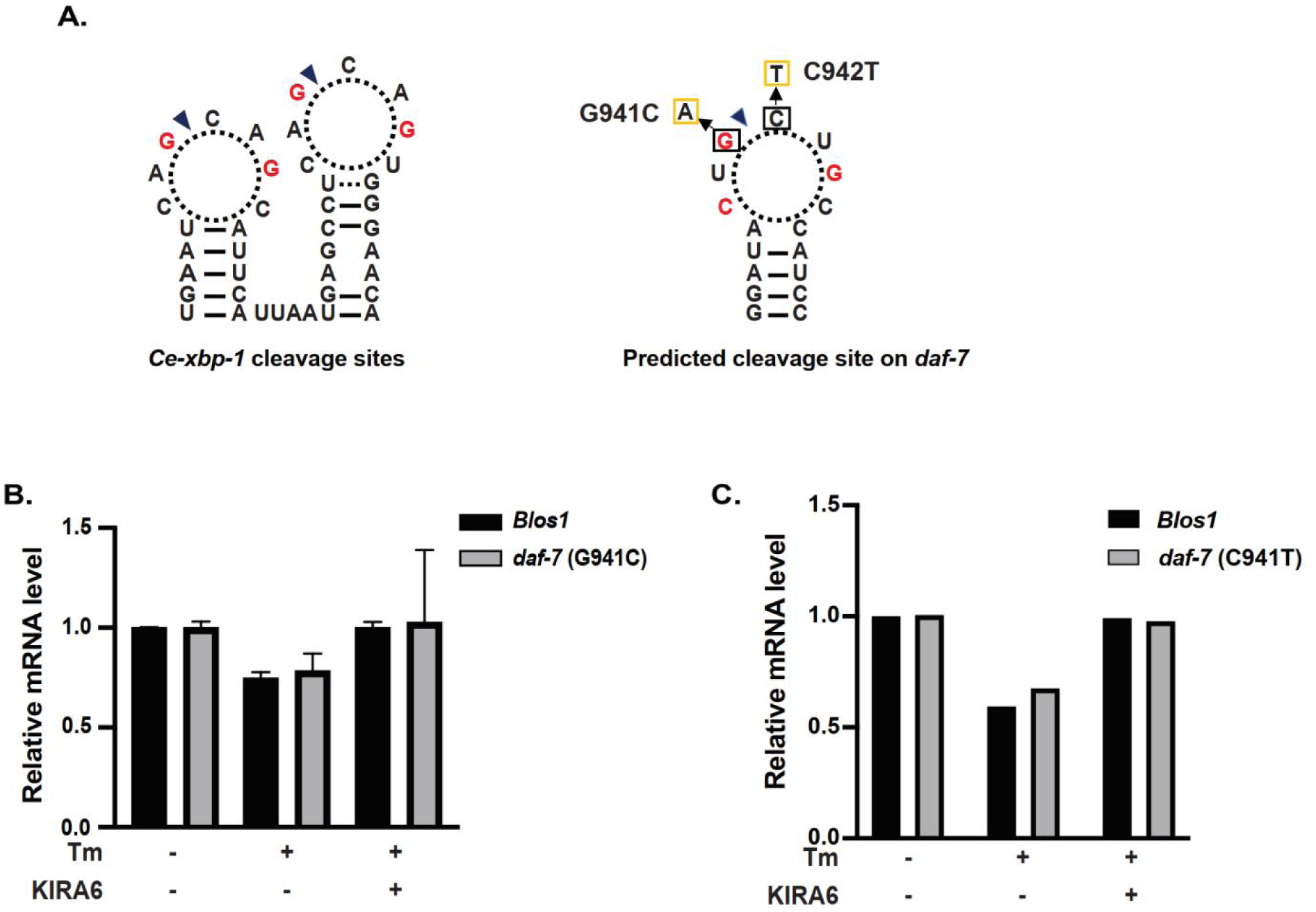
*daf-7* mRNA likely contains IRE1 cleavage sites that are different from the canonical one. Related to Figure 3. **(A)** Predicted IRE1 recognition and cleavage sites on *daf-7* mRNA. Left, double-stem loop and cleavage site(s) (arrowheads) in the known *C. elegans* IRE1 substrate *xbp-1* mRNA; right, stem loop with a predicted recognition and cleavage (arrowhead) site on *daf-7* mRNA. The potential cleavage site (dark triangle) is shown on the right, mutations are indicated. (**B and C)** Mutating predicted IRE1 cleavage site does not affect *daf-7* downregulation. RT-qPCR analysis of *Blos1* and mutated *daf-7* mRNA in WT HAP1. Cells treated as in Fig. S4. 3 independent experiments for *daf-7* with G941C mutation, error bars: mean+/-SD.

**Figure S6.**
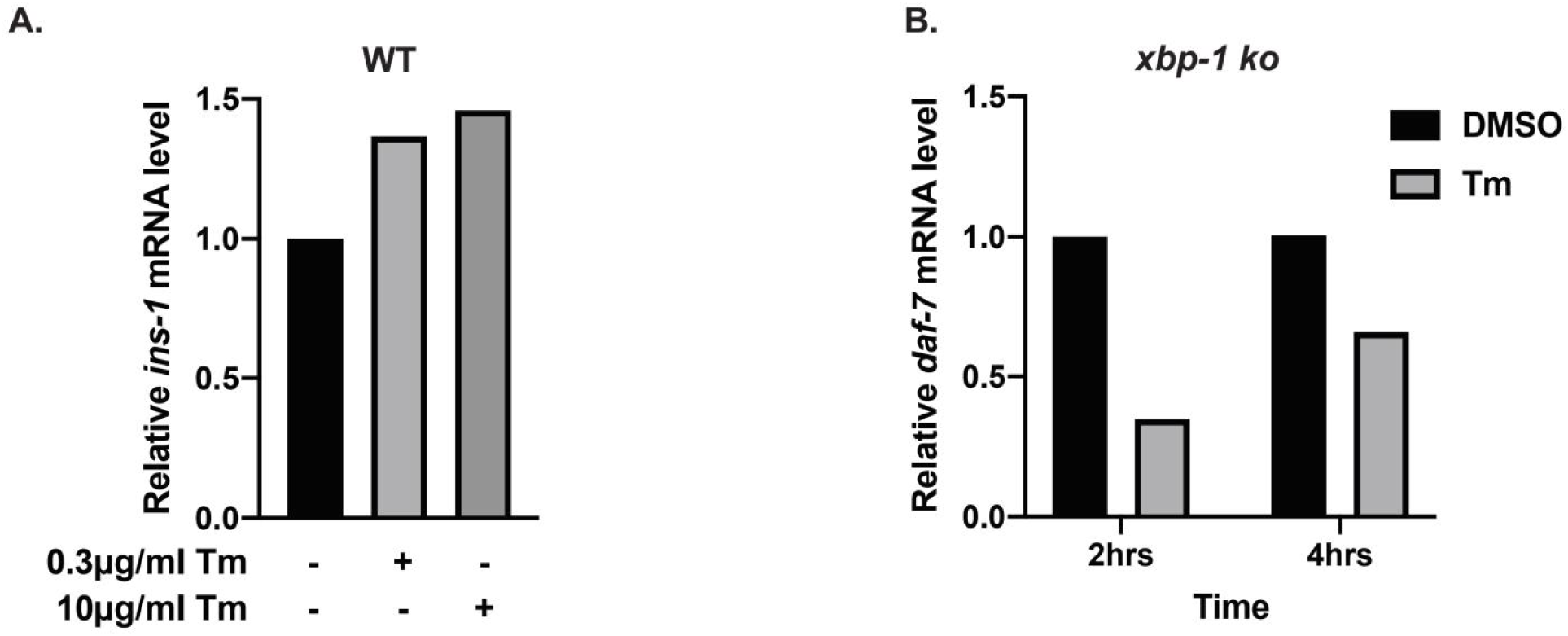
RIDD in *C. elegans* is XBP-1 independent and substrate selective. Related to Figure **(A)** *ins-1* mRNA is not downregulated by IRE1 in *C. elegans.* Late L1 larvae were treated with either 0.3μg/ml or 10μg/ml Tm for 4hrs and analyzed by RT-qPCR. The *ins-1* transcript level change was normalized to *actin*. **(B)** Downregulation of *daf-7* mRNA by activated IRE1 is XBP-1-independent. Late L1 animals were treated with 10μg/ml Tm for 2hrs or 4hrs. *daf-7* transcripts were normalized to *actin*.

**Figure S7.**
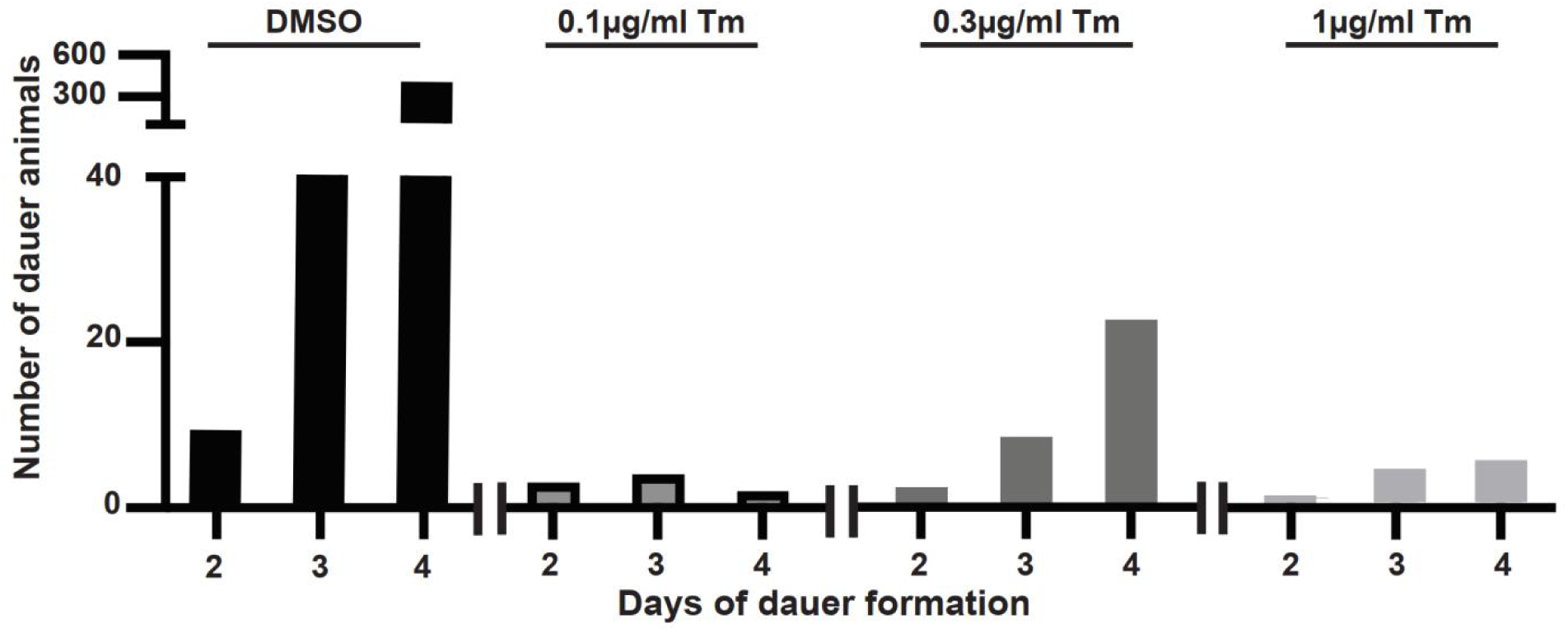
Tm treatments inhibit dauer transition even at low concentrations. Related to Figure 5. Two wild type L4 animals were placed onto small plates freshly seeded with bacteria and containing indicated concentration of Tm at 25°C. The animals were allowed to grow until all food was exhausted, and number of dauers was determined after 2, 3 and 4 days of starvation.

**Figure S8.**
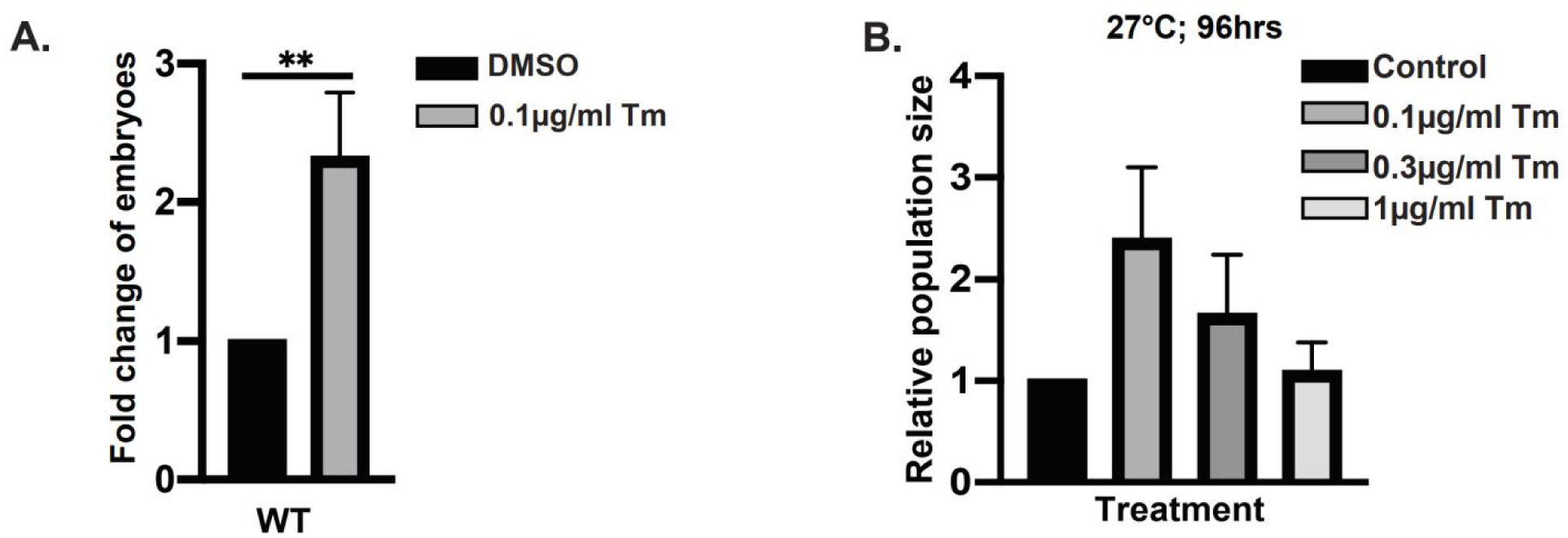
Pre-treatment with low Tm concentrations improves reproduction and population growth under environmental stress. Related to Figure 5. (**A)** Wild type embryos were plated as in Figure 5A and grown at 27°C with or without 0.1μg/ml Tm. After 72hrs, the number of embryos present on plates was counted. 3 independent experiments, error bars are means with SD, *t-*test, significance as described in material and method. **(B)** Low Tm concentrations are still protective when animals are allowed to grow at 27°C for extended times, reaching complete starvation 96hrs. Plates were set up as in Figure 5A, the data shown are from 3 independent experiments.

**Figure S9.**
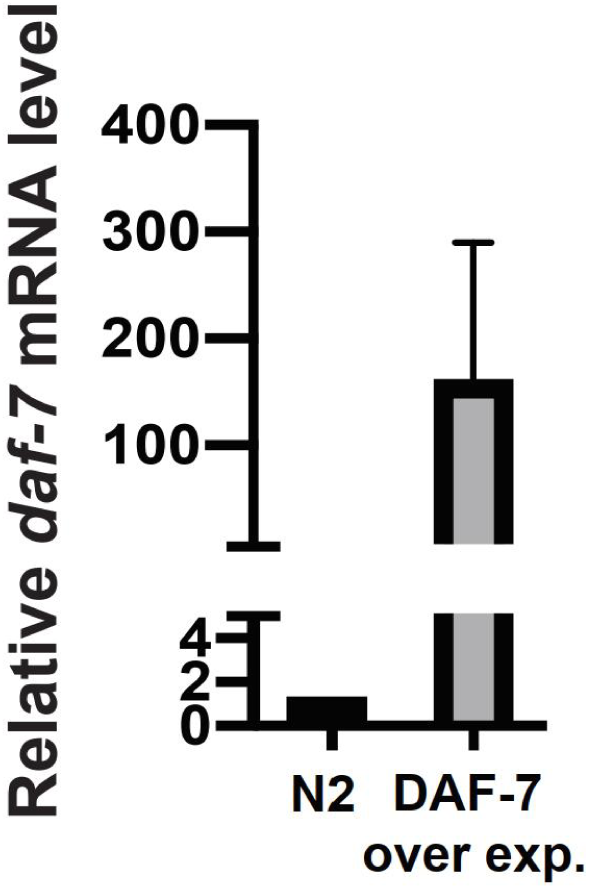
Quantitation of *daf-7* overexpression in transgenic animals. Related to Figure 5. RT-qPCR analysis of *daf-7* mRNA level in late L1 larvae, normalized to actin. DAF-7 over exp.: p*daf-7::mcherry::daf-7*) . 3 independent experiments, error bars: mean+/-SD.

## Material and methods

### Cell culture and viral infection

HAP1 IRE1α KO (HAP1 IRE1KO) is an engineered HAP1 cells (a near-haploid human cell line) with 8 base pairs deleted in exon 6 of endogenous IRE1α using CRISPR-Cas9 were purchased from Horizon (HZGHC000742c006). HAP1 and HAP1 IRE1KO cells were cultured in Iscove’s Modified Dulbecco’s Medium (IMDM) (Invitrogen) with additional 5% fetal bovine serum (Atlanta Biologicals, Flowery Branch, GA, USA), 1% sodium pyruvate (Corning, Corning, NY, USA) and 1% penicillin/streptomycin (Thermo Fisher Scientific).

HAP1 IRE1KO cells were infected with a pTight Tet-On lentiviruses coding for either wild type or mutant IRE1 chimeras, or mCherry-DAF-7. pTight lentiviral vector with human IRE1-GFP was used as the backbone^58^. Infected HAP1 cells were selected with puromycin, and IRE1 clones were chosen based on the protein expression to avoid overexpression, since it can alone activate IRE1. Expression of transgenes was induced with 1μg/ml doxycycline (dox) (Biochemika) for 18 hours. Tunicamycin, 4μ8C, KIRA6 were from Calbiochem. Thapsigargin was from MP Biomedicals, and Brefeldin A were from MilliporeSigma.

### Transfection

HAP1 IRE1KO expressing human IRE1-GFP cells were transfected with pBud4 plasmid carrying full length (unspliced) *C. elegans xbp-1.* Transfections were done using Lipofectmine 2000 (Invitrogen), for 6hrs. Cells were incubated overnight, trypsinized, and split into in 6-well plates for pharmacological treatments.

### Mutagenesis

DAF-7-mCherry lentiviral plasmid was used as template to introduce G941C or C942T mutations (nucleotide numbers based on cDNA). Site-directed mutagenesis was performed with Pfu Ultra II Fusion HS polymerase (Agilent). Mutations were confirmed with Sanger sequencing.

### Selection of candidate RIDD targets

We used data from microarray analyses in Shen *et al.*, 2005^51^, which determined transcriptional changes upon Tm treatment in L2 animals of wild type strain and strains with individually deactivated IRE1, XBP1, PERK and ATF6. 43 candidate transcripts from Table 5 in Shen *et al.* (3538 transcripts total) were selected in the following way: we first chose genes that were downregulated by Tm treatment by at least 25% in the wild type (N2) animals. From these, we next selected transcripts whose downregulation depended on IRE1 ((ΔN2-ΔIRE1KO)/ΔN2≥0.9), resulting in 112 transcripts. Finally we selected from the remaining transcript those that showed no or low dependence on XBP1 (with threshold of 20% downregulation) and PERK or ATF6 (10 % downregulation). Removing duplicates and genes without functional annotation resulted in 43 candidate genes, 30 of which are either know or predicted to be coding for secretory proteins (Table 1).

### C. elegans strains

Nematodes were grown at 20°C on nematode growth medium (NGM) plates seeded with *Escherichia. coli* OP50^108^. Animals were synchronized as described for tunicamycin treatment below. Wild type *C. elegans* strain (Bristol N2) and the following strains: SJ4005(*zcIs4[*p*hsp-4*::GFP]), SJ30(*ire-1(zc14)* II), zcIs4[p*hsp-4::*GFP]), RB545(*pek-1(ok275)* X), SJ17(*xbp-1*(*zc12*) III) were obtained from the Caenorhabditis Genetic Center. DAF-7 overexpression strain drxEx20 [p*daf-7::mCherry::daf-7::*unc-54 3’UTR] is described in ref^72^.

### Activation of *C. elegans* IRE1 by tunicamycin treatment

Care has to be taken throughout the described steps that animals are not crowded and have plentiful food, as even small amounts of nutritional or density stresses affect UPR induction. ∼500 embryos were collected from well maintained (for at least three generations) *C. elegans* plates by scraping, to generate first roughly synchronized population. This population was allowed to grow on fresh 10cm plates for 96 hours, to reach the maximum egg-laying rate. Second roughly synchronized population was generated by washing off larvae and adults, collecting embryos by scraping, and transferring them to new NGM plates. After incubation at 20°C for 96hrs, adults were transferred to fresh NGM plates and allowed to lay eggs for 90 mins (∼1000 eggs were laid). For tunicamycin treatments, embryos were allowed to hatch and grow for 28hrs. The hatched (fine synchronized) young larvae then were transferred to assay plates containing defined concentrations of drugs, as needed.

### Tm-containing plates for *C. elegans* treatment

To prepare Tm-containing plates, Tm stock (10mg/ml in DMSO) was first diluted in M9 buffer (1/200 of total medium for NGM plate) supplemented with 1μl of 3M NaOH to prevent precipitation. This solution of Tm was added to warm NGM-agar solution (65-70°C) to the desired concentration under rapid mixing.

### RT-qPCR in human cells

Total RNA was extracted with Trizol (Life Technologies) following manufacturers protocol. 200ng of extracted RNA was retro-transcribed to cDNA by using oligo (dT)12-18 (Invitrogen) and SuperScript II reverse transcriptase (Invitrogen). Quantitative PCR was performed using SYBR Green reagent (Applied Biosystems) and Applied Biosystems StepOne Plus thermocycler with annealing temperature of 60°C, primer sequences are provided below. Data were analyzed by normalizing to *Rpl19* using the ΔΔ-Ct method.

### RT-qPCR in *C. elegans*

Total RNA was extracted from ∼1000 synchronized late L1 animals using Trizol (Life Technologies) following manufacturers standard protocol. 100ng total RNA was converted to cDNA, and analyzed as described for RT-qPCR in human cells. Data were analyzed by normalizing to *actin* using the ΔΔCt method.

### Western blots

HAP1 cells were lysed using following lysis buffer: 50 mM Tris-HCl pH 8, 150 mM NaCl, 5 mM KCl, 5 mM MgCl2, 1% NP-40, 1X complete protease inhibitors (Roche). Protein concentration was determined by BCA assay (Thermo Fisher). Lysates were resolved on 7 or 10% gels by SDS-PAGE, and transferred onto nitrocellulose membranes (Bio-Rad). Membranes were blocked with milk, probed with primary and secondary antibodies, and scanned with Odyssey Infrared imager (Li-Cor). Primary antibodies used were: anti-N term IRE1 (GeneTex); anti-mCherry (Thermo Fisher Scientific); anti-α tubulin (Sigma). IRDye-conjugated secondary antibodies were from Li-Cor.

### *daf-7* RNA stem-loop structure prediction

Stem-loop structure was predicted using ViennaRNA Web Services (http://rna.tbi.univie.ac.at/) with minimum free energy (MFE) model. Short RNA sequences (around 30 base pairs) were used as input for the algorithms.

### Population growth assay

25 embryos were seeded onto Tm-containing 3cm NGM plates with plentiful food. Embryos were incubated at 27°C or 20°C for 72 hours or 96 hours, the number of offspring larvae were counted in each condition by sampling, and normalized to the number of larvae in NGM plates without Tm.

### Dauer induction assay

L4 larvae were transferred to peptone-free NGM plates (two per plate) with concentrated OP50 bacteria, and incubated at 25°C until plates were starved. Dauer animals were counted after 48hrs, 72hrs or 96hrs. Dauers were identified by the resistance to 1% SDS.

### Imaging

Confocal: Transfected cells were imaged in μ-Dish 35mm culture plates (Ibidi) with Zeiss LSM700 microscope equipped with CO2 and temperature-controlled chamber, at Cell Imaging Center, Drexel University. Cells were imaged with 1.4NA 63x oil objective, and 12 bit confocal stacks were reconstructed in ImageJ as projections.

Stereo: animals were immobilized with sodium azide on chilled unseeded NGM plates and imaged with Leica M205FA microscope and Hamamatsu Orca R2 camera, keeping magnification and intensity of fluorescent sources (Chroma PhotoFluor 2) constant within experiment. All animals within one experiment were imaged in one session. Fluorescence intensity was quantified with ImageJ.

### Statistics

Unpaired *t-test* analyses were performed using Prism software (GraphPad, USA). significance levels are indicated in the assays with at least three repeats.

Data were analyzed with *t*-test or one-way ANOVA with Tukey multiple comparison correction. All analyses were performed using Prism software (GraphPad, USA). α value in all experiemnts was 0.05, and significance levels as follows: ns: P > 0.05; *: P ≤ 0.05; **: P ≤ 0.01; ***: P ≤ 0.001; **** means P ≤ 0.0001.

### List of primers used

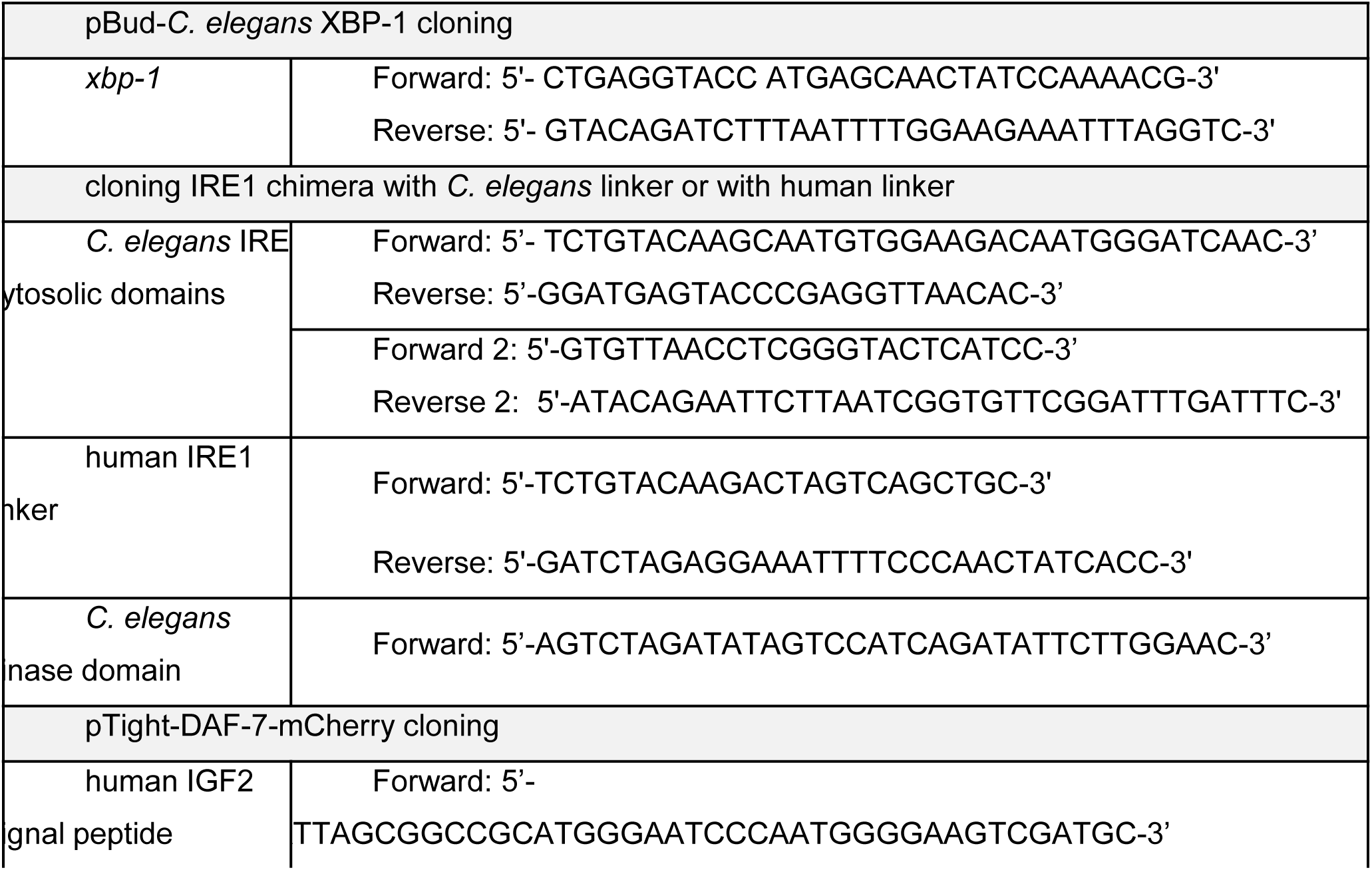
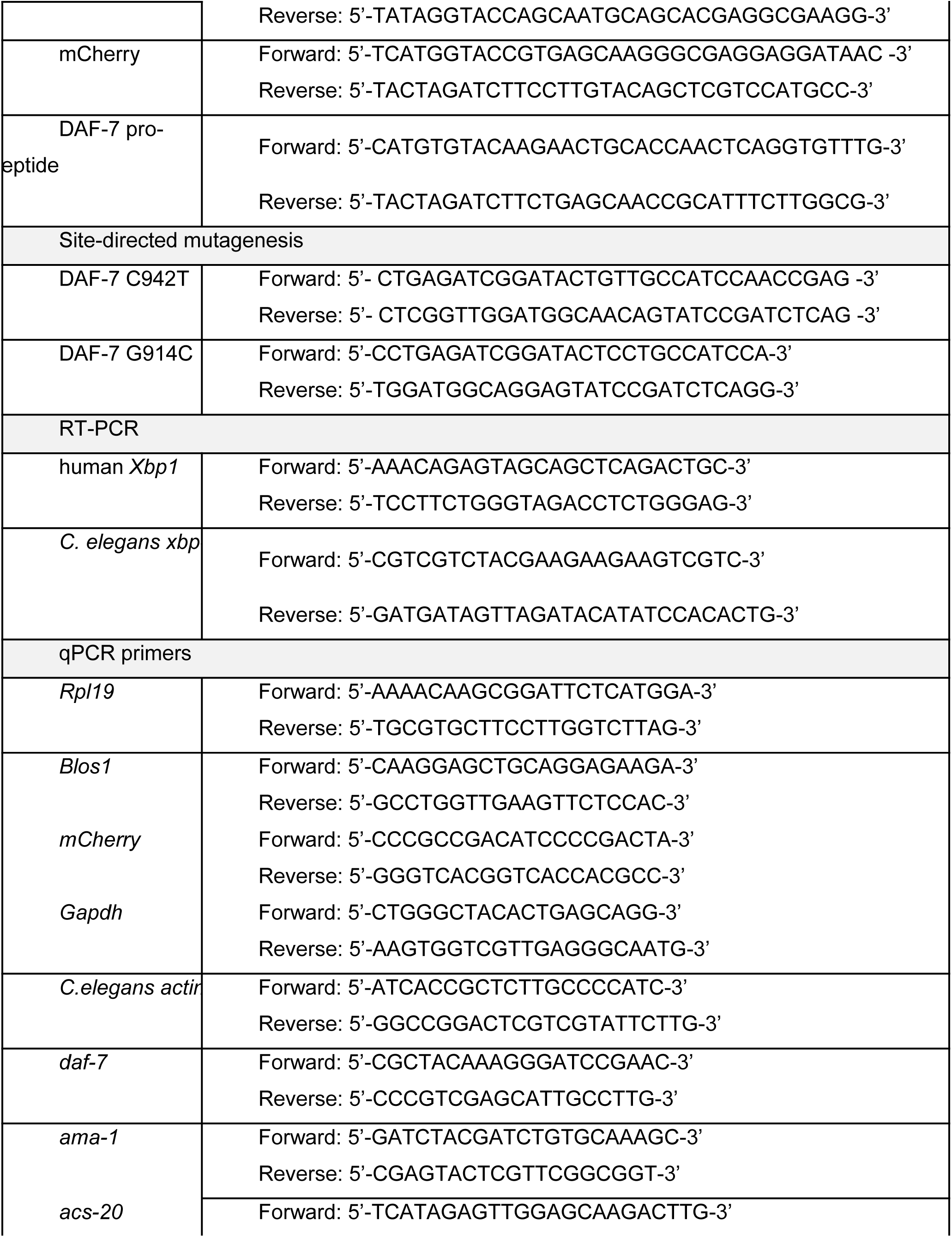
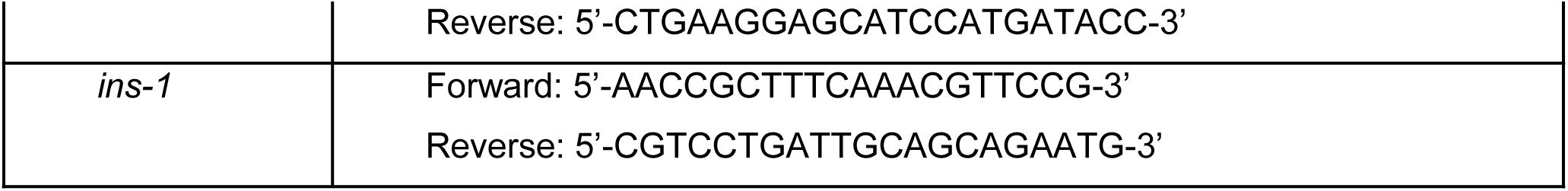

